# Cryo-EM inspired NMR analysis reveals a pH-induced conformational switching mechanism for imparting dynamics to Betanodavirus protrusions

**DOI:** 10.1101/2024.03.08.584019

**Authors:** Petra Štěrbová, Chun-Hsiung Wang, Kathleen J.D. Carillo, Yuan-Chao Lou, Takayuki Kato, Keiichi Namba, Der-Lii M. Tzou, Wei-Hau Chang

## Abstract

Nervous necrosis virus (NNV), a non-enveloped betanodavirus, causes neuropathies and retinopathies in farmed fish, damaging aquaculture worldwide. NNV has 60 conspicuous surface protrusions comprising the protrusion domain (P-domain) of its capsid protein. Although NNV protrusions play critical roles in infectivity, the underlying dynamics remain unclear. Our cryogenic electron microscopy (cryo-EM)-derived structures of Dragon grouper (*Epinephelus lanceolatus*) NNV reveal that the protrusions undergo low-pH-induced compaction and movement. We show that the P-domain is monomeric in solution at a pH germane to infection (7.0). Moreover, nuclear magnetic resonance (NMR) structures reveal a peptide (amino acids 311-330) that adopts a flexible loop to form an open pocket. NMR spectral analysis at pH 5.0 aided by molecular dynamics (MD) simulations show that this loop switches to a β-strand under acidic conditions, eliciting pocket closure and P-domain trimerization, highlighting a unique pH-sensing feature. Our docking analysis revealed the N-terminal moiety of sialic acid inserted into and interacting with conserved residues in the pocket. Additionally, a low-pH-induced conformational change in the linker region via peptide bond isomerization conferred malleability on the protrusions. Our work uncovers the protrusion dynamics of a betanodavirus governing its infectivity through a pH-dependent conformational switching mechanism, providing insights into complex virus-host interactions.

## Introduction

Nervous necrosis virus (NNV) has become a primary infection threat for industrial fish farming worldwide [1,2]. Outbreaks of the viral neuropathies and retinopathies caused by NNV result in almost 100% mortality of infected stock, with larval and juvenile fish being the most susceptible to NNV infection. Since first characterization of the virus from larval striped jack (*Pseudocaranx dentex*) in the 1990s [3], NNV infections have been reported in more than 120 marine and freshwater fish [4]. Due to the lack of a vaccine, NNV infections continue to damage significantly the aquacultural economy.

NNV is a simple spherical non-enveloped RNA virus of Genus *Betanodavirus* and Family *Nodaviridae*. The NNV viral particle encapsulates two positive-sense single-stranded RNA molecules, RNA1 and RNA2. The larger RNA1 (3.1 kilobases, kb) encodes RNA-dependent RNA polymerase, whereas the smaller RNA2 (1.4 kb) encodes a single structural capsid protein (CP), with an apparent molecular mass of 42 kDa [5]. A sub-genomic RNA (RNA3), comprising part of the 3’ end of RNA1, has been identified as encoding non-structural protein B2, which acts as an antagonist to host RNA interference during infection [6]. There are two genera in Family *Nodaviridae*, i.e., alphanodaviruses that primarily infect insects, and betanodaviruses whose natural hosts are fish [5]. However, the CP sequences of alphanodaviruses and betanodaviruses share low similarity, with alphanodaviruses lacking the pronounced surface protrusions of betanodaviruses [7].

NNV has been demonstrated to enter host cells via an endocytic pathway [8–10]. During this process, viruses are internalized into endosomes for destruction. However, the highly acidic conditions inside endosomes (pH of 5.0–5.5) can activate release of the viral genome [11–13], perhaps by altering the capsid structure, as exemplified by the non-enveloped alphanodavirus Flock House Virus (FHV). Exposure of FHV to low endocytic pH induces structural changes in the FHV viral particle and release of its membrane-interacting γ peptide (4.4 kDa) that is capable of disrupting the endosomal membrane [14–15]. However, the impact of acidic pH on betanodavirus particles during endocytosis remains unclear.

An atomic model of the NNV capsid was obtained previously based on X-ray diffraction of crystals of Grouper NNV (GNNV) virus-like particles (VLPs) to near atomic resolution (3.6 Å, PDB 4WIZ) [16]. This X-ray structure, obtained under neutral conditions (pH 6.5), proved consistent with a previously generated low-resolution cryo-EM structure [7], which unveiled the GNNV capsid as being an icosahedral particle (T=3) with a diameter of ∼35 nm. Moreover, the X-ray struture resolved the CP as having three domains: an N-terminal arm, a shell domain (S-domain), and a protrusion domain (P-domain). The positively-charged flexible N-terminal arm is located inside the viral particle to assist in viral RNA packing during particle assembly. The S-domain exhibits a canonical “jelly-roll” topology comprising an eight-stranded β-sheet for building the virus capsid. The P-domain is located outside of the viral particle and is connected to the S-domain via a flexible linker. The P-domains from three contiguous CPs in one asymmetric unit form a protrusion at the quasi-three-fold axis. These protrusions play a critical role in NNV infectivity and antigenicity [17,18], with the variable region of the P-domain [19] determining NNV host specificity [20, 21]. The P-domain has been crystallized separately and resolved to atomic resolution (1.2 Å, PDB 4RFU) [16]. In that crystal structure, three P-domain molecules self-assemble into a non-crystallographic trimer, which is stabilized by two calcium ions coordinated by aspartic acid residues on the trimer interface. Those crystal structures [16] provide a static view for GNNV while potential dynamics of protrusions still remain unknown.

Compared to X-ray crystallography, cryo-EM represents an alternative high-resolution technique with the advantages of providing easier access to close-to-native state structures in conditions mimicking native environments and allowing for co-existing conformations to be disentangled [22]. Those advantages are crucial for capturing snapshots of dynamic structures or structural alterations induced by changes in pH, ions [23], or temperature [24]. Herein, we used cryo-EM imaging to explore structural alterations of GNNV particles in response to pH changes, allowing us to determine the structures of native GNNV virions (pH 6.5 and 5.0) and virus-like particles (pH 8.0, 6.5 and 5.0) to near atomic resolutions (2.82 to 4.36 Å). Our results reveal that the protrusions on GNNV became erect under conditions suitable for GNNV host cell entry (pH 8.0-7.0), but rested on the capsid under the acidic conditions encountered in late endosomes (pH 5.0). To further resolve the dynamics of the protrusions, we expressed the GNNV P-domain separately and analyzed its solution behavior to show that it is a monomer at pH 7.0, but could convert to a trimer at pH 5.0. Further NMR-derived structural determinations of the P-domain (pH 7.0) yielded its solution structure, which uncovered a long peptide comprising amino acids (aa) 311-330 that adopts a flexible loop, contrasting with the secondary structure in the crystal structure [16]. We used this solution structure as the basis for molecular dynamics (MD) simulations to investigate pH-dependent structural transitions, leading to identification of a region within the flexible loop that may acquire secondary structure at low pH to allow for P-domain trimerization. These findings uncover a detailed mechanism underlying the dynamics of GNNV protrusions, providing an authentic structural basis for investigating the mechanism of betanodavirus infectivity and for designing a potential vaccine.

## Results

### Cryo-EM reveals that GNNV protrusions are dynamic entities

Close inspection of a high-quality cryo-EM map of a GNNV VLP generated previously (see **Fig. 1B** in [25]) against the respective crystal structure (see **Fig. 1D** in [16]) suggests that the protrusions in solution state are morphologically less compact. However, it is unclear whether this discrepancy is attributable to crystal packing or due to differences in the buffer conditions employed in these two studies (pH 6.5 for crystal [16] and pH 8.0 for the cryo-EM [25]). Therefore, we set out to investigate potential GNNV structural changes in different pH environments using multi-purposed cryo-EM enabled by a direct electron detection camera [25]. We immersed native GNNV virions or respective VLPs in buffers of three different pH, i.e., 8.0, 6.5, and 5.0, corresponding to seawater, GNNV VLP crystallization conditions [16], and the acidic environment inside late endosomes, respectively. Interestingly, our cryo-EM images revealed a spikier appearance of the GNNV VLPs at pH 8.0 and pH 6.5 than at pH 5.0 (**Fig. S1**). By using 3D reconstruction and imposing icosahedral symmetry, we determined cryo-EM structures for GNNV VLPs at pH 8.0, 6.5 and 5.0 (**Fig. 1**), as well as for virions at pH 6.5 and 5.0 (**Fig. S2**), with overall resolutions in the range of 2.82 Å to 4.36 Å **(Fig. S3** & **S4)**. These results indicate that lowering the pH prompted GNNV particles to undergo gross structural alteration (**Fig. 1** & **Fig. S2**), with VLP and virion structures at the same pH being virtually indistinguishable (**Fig. S2**), validating that VLPs represent good surrogates of GNNV virions for vaccine development [26].

**Figure 1.**
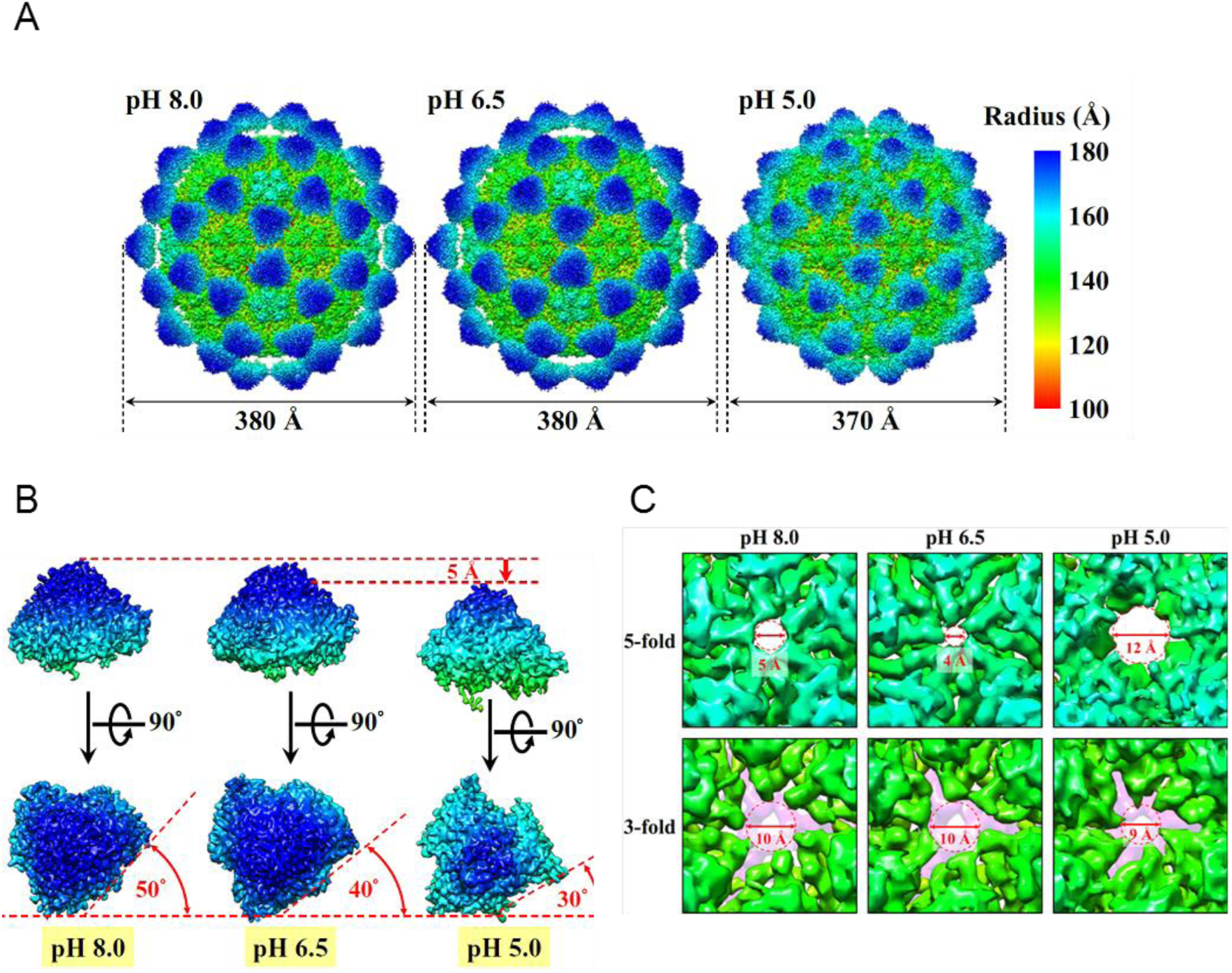
Cryo-EM structures of GNNV in three different pH environments. **(A)** Surface views of GNNV virus-like particles at pH 8.0 (*left panel*), pH 6.5 (*middle panel*) and pH 5.0 (*right panel*). The views are colored according to the local radius heatmap shown at right. **(B)** Conformational changes of the P-domain upon switching from pH 8.0 (*left panel*) to pH 6.5 (*middle panel*) and pH 5.0 (*right panel*). The protrusions are rotated clockwise and shift ∼5 Å toward the capsid shell upon changing the pH. **(C)** Enlarged views (5-fold and 3-fold) of pores in the capsid shell surface for different pH environments. The size of each pore for each pH condition is indicated. In order to better reveal the pore at the 3-fold axis, the surface densities of the underlying N-terminal domains of the capsid protein (CP) have been colored light purple.

Interestingly, the diameter of the GNNV particle in the acidic condition (pH 5.0) was smaller (370 Å) than under alkaline (pH 8.0) or neutral (pH 6.5) conditions (380 Å) (**Fig. 1A**). This reduced size is due to a ≈5Å displacement of the protrusions toward the capsid shell (**Fig. 1B**), with radial density and cross-section analyses showing that shell size remained virtually unchanged (**Fig. S5** & **S6**). Moreover, shell morphology was relatively unperturbed by the acidic condition (**Fig. 1C**). Closer examination of the protrusions revealed that not only were they positioned closer to the shell in the acidic pH, but they also became more compact and were rotated clockwise by ∼20° (**Fig. 1B, Movie S1** & **S2**). This repositioning prompted our hypothesis that the linker between the P-domain and S-domain is malleable (**Movie S3**), prompting further investigation of the detailed mechanism (**Fig. S3** & **S4**). In sum, our cryo-EM imaging has captured the pH-responsive structures of native GNNV virions, revealing that the capsid protrusions in solution adopt an erect position under alkaline and neutral conditions, but a prone position under acidic conditions.

### NMR spectra indicate that the P-domain of GNNV contains highly pH-sensitive regions

To investigate the behavior in solution of the protrusions as isolated entities instead of on the GNNV capsid, we cloned and expressed the P-domain (aa 221-338) plus the flexible linker (aa 214-220, linking the P-domain and S-domain) from GNNV (GNNV-P) for structural analysis using NMR spectroscopy and sedimentation velocity analytical ultracentrifugation (AUC). GNNV-P has a molecular weight of 13.8 kDa. Notably, neither the N-terminal linker (aa 214-219) nor C-terminal region (aa 337-338) were observed in a previously generated X-ray structure of GNNV-P (PDB 4RFU), likely due to their high flexibility [16]. In the buffer we used for solution analyses, we excluded divalent ions or small molecules used to induce the crystal [16].

We previously collected 2D ^1^H–^15^N HSQC spectra for GNNV-P at pH 7.0 [27]. To explore the pH-dependence of GNNV-P, we conducted a pH titration experiment to cover a wide range of pH values (7.0, 6.0, 5.8, 5.5, 5.2, and 5.0; see **Fig. 2A** for pH 7.0 and pH 5.0). Our pH titration experiment revealed signatures of pH-sensitive residues, the peaks of which disappeared as pH decreased, though some reappeared at different positions (**Fig. 2B**). Moreover, we observed a significant line broadening effect at pH 5.0 (**Fig. 2A**), indicating that GNNV-P may undergo a low-pH-induced conformational change and/or oligomerization. To test if this pH-dependent property is reversible, we monitored the 2D HSQC spectra by titrating at increasing pH from 5.0 to 7.0, which showed that the pattern for each pH value could be reproduced, indicating that the pH-dependent changes are reversible (data not shown).

**Figure 2.**
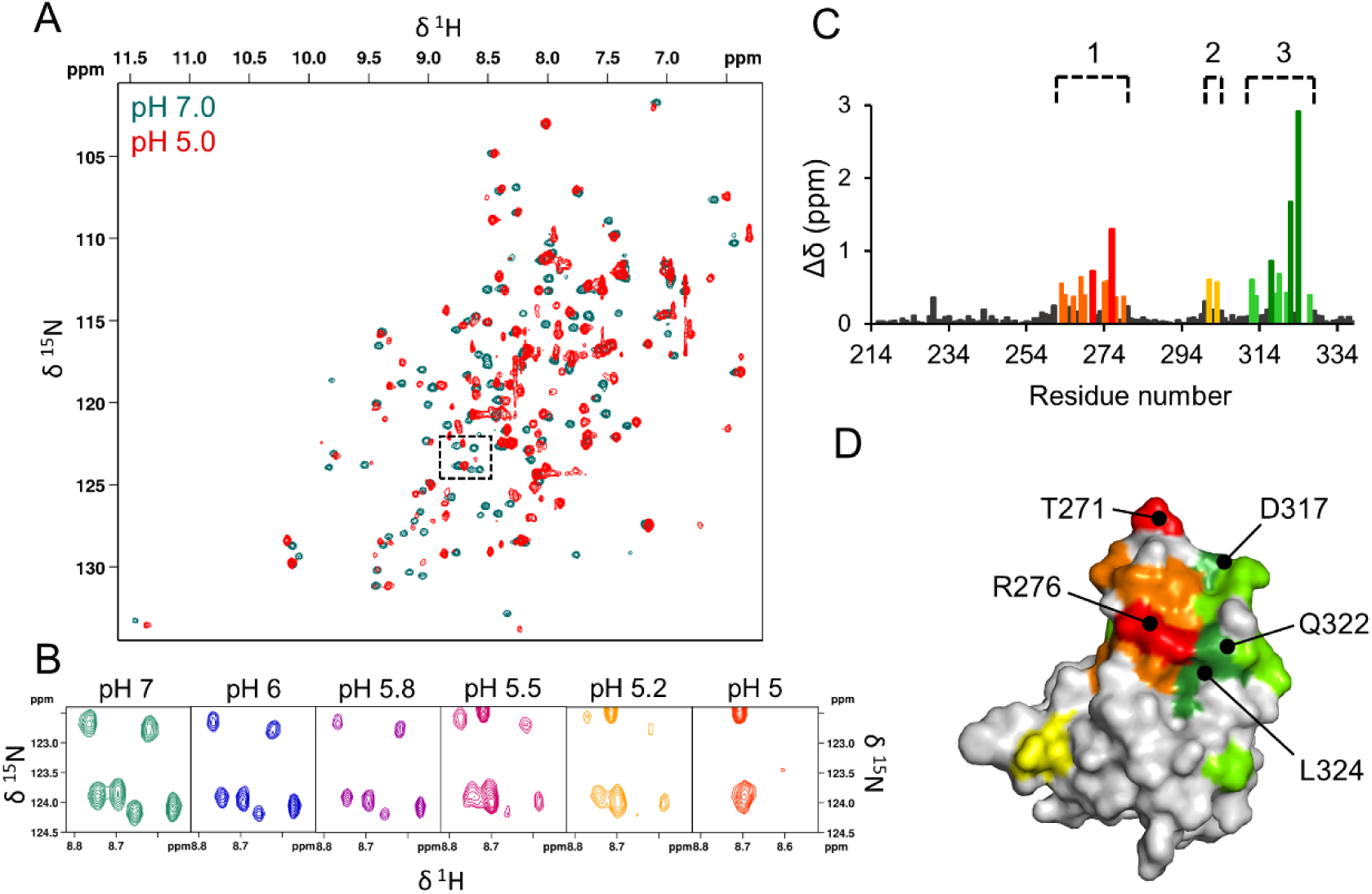
Effects of pH on GNNV-P in solution. **(A)** ^15^N HSQC spectra of GNNV-P recorded at pH 7.0 (blue) and pH 5.0 (red). **(B)** Detail of the boxed area of the ^1^H-^15^N HSQC spectra in (A), showing the changing peak pattern during pH titration of GNNV-P from pH 7.0 to pH 5.0. **(C)** Summary of chemical shift perturbations (CSPs) due to changing from neutral to acidic pH. The pH-sensitive regions I (L263-Y279), II (W301-N303), and III (V312-V327) are highlighted in orange, yellow, and green, respectively. Residues with CSPs >2 standard deviations in pH-sensitive Region I and Region III are highlighted in red and dark green, respectively. **(D)** pH-sensitive residues mapped onto a surface representation of the GNNV-P crystal structure [16], with the same color scheme as in (B).

To improve the quality of the pH 5.0 spectra, we utilized deuterium labeling since replacing the ^1^H nuclei with D (^2^H) can suppress strong ^1^H-^1^H dipolar interactions. The resulting uniformly ^13^C-, ^15^N-, and D-labeled GNNV-P generated a well-resolved 2D HSQC pattern at pH 5.0 (**Fig. S7**), allowing us to assign backbone resonances. We identified 98.3% of the expected amide ^1^H_N_ and ^15^N_H_ resonances (115 out of 117 non-proline residues), with the exception of residues T214 and L325 (both of which precede prolines in the sequence). Moreover, backbone ^13^C_α_, ^13^C_β_, and ^13^C’ resonances were assigned with 98.2-98.4% completeness (see **Data Availability**).

Chemical shift perturbations (CSPs) reflect changes in the chemical environment of atomic nuclei. These changes can arise from protein interactions or conformational changes. Mapping residues displaying CSPs onto a protein structure can help identify interaction sites [28]. In principle, CSPs can be analyzed for each residue at each pH titration point (see **Fig. 2B**), but it is challenging to track chemical shifts that disappear and then re-appear at positions differing substantially from the original ones. We thus opted to analyze GNNV-P CSPs by comparing only the chemical shifts at pH 7.0 and 5.0, i.e., the initial and final points of our pH titration experiments (**Fig. S8**). At pH 5.0, most signals had re-appeared (**Fig. 2A** & **Fig. S8**). With nearly all of the chemical shifts assigned for the ^1^H_N_ and ^15^N_H_ data at pH 7.0 [27] and 5.0 (**Fig. S7**), we applied CSPs analysis [29] to identify residues displaying significantly perturbed chemical shifts relative to pH 7.0, i.e., CSPs greater than one standard deviation from the mean. In **Figure 2C**, we show the backbone CSPs per residue based on amide nitrogen and proton chemical shifts. Intriguingly, residues displaying pronounced CSPs are clustered into three regions: Region I (L263-Y279), Region II (W301-N303), and Region III (V312-V327). Regions I and III are spatially proximal to each other, as determined by mapping the pH-sensitive residues onto the GNNV-P crystal structure [16] (**Fig. 2D**). Since residues with pronounced CSPs are likely involved in protein interactions and/or conformational changes [28], we speculate that these three regions undergo low-pH-induced interactions or conformational changes. Notably, these regions in GNNV-P that confer pH sensitivity coincide with three β-strands in the crystal structure [16].

### Determination of the structure of GNNV-P in solution at pH 7.0 reveals that aa 311-330 form a long flexible loop

Based on the X-ray crystal structure determined under neutral conditions [16], GNNV-P was assumed to form a trimer in solution at pH 7.0. Surprisingly, our AUC experiments showed that GNNV-P molecules predominantly adopted a monomeric form at pH 7.0, and only assumed the trimeric form in an acidic condition (pH 5.0) (**Fig. 3A** & **Fig. S9**). Accordingly, we set out to determine the solution structure of GNNV-P based on the HSQC spectra at pH 7.0 (**Fig. 2A**), which exhibited well-dispersed cross-peaks with sharp resonances, indicative of a well-folded GNNV-P structure in solution.

**Figure 3.**
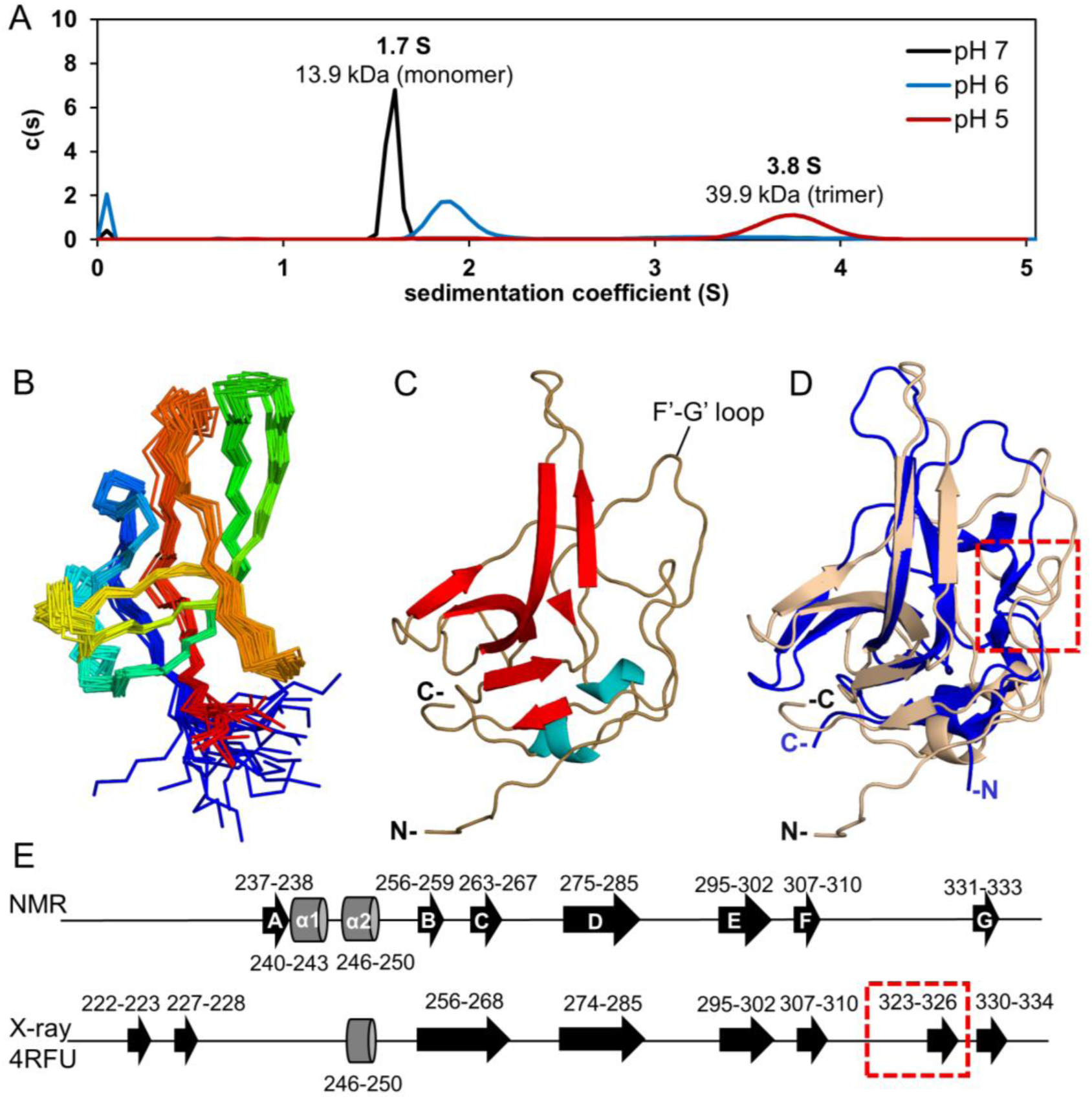
Structure of GNNV-P determined in solution at neutral pH. **(A)** Sedimentation velocity analytical ultracentrifugation (SV AUC) data of GNNV-P obtained at pH 7.0 (black), pH 6.0 (blue), and pH 5.0 (red) fitted to a continuous sedimentation coefficient distribution c(s) model. Sedimentation coefficients and the molecular weight determined by SEDFIT are noted above each peak. **(B)** Backbone (N, Cα, C’) superimposition of the ensemble of 20 low-energy conformations. The N-terminal is colored blue and the C-terminal is in red. **(C)** Cartoon representation of the GNNV-P solution structure with β-strands and α-helices highlighted in red and blue, respectively. **(D)** Superimposition of the GNNV-P structure determined by NMR at neutral pH (pink) and by X-ray crystallography (PDB 4RFU, blue).. **(E)** Schematic representation of NNV P-domain secondary structures, as determined by NMR (pH 7.0) and X-ray crystallography (pH 6.5). β-strands are displayed as black arrows and α-helices as grey barrels, with bordering residues indicated.

From the ^1^H, ^13^C, and ^15^N backbone and side-chain chemical shift assignments for 122 GNNV-P residues (except for Thr214, Leu228, and Gly270) at pH 7.0 [27], we calculated the solution structure of GNNV-P using nuclear Overhauser effect (NOE)-derived ^1^H-^1^H distance restraints, chemical shift-derived dihedral angle restraints, and hydrogen bonds inferred from hydrogen deuterium exchange (**Fig. S10**) (see **Materials and Methods** for details of NMR structural determination). The respective calculation statistics are summarized in **Table 2**.

In **Fig. 3B**, we present an ensemble of 20 low-energy solution structures, with a representative structure illustrated in **Fig. 3C**. This solution structure of GNNV-P is cone-shaped, similar to the crystal structure (PDB 4RFU; see superimposed structures in **Fig. 3D**). The folded GNNV-P tertiary structure comprises a number of secondary structures including anti-parallel β-strands and α-helixes, with the order βA-α1-α2-βB-βC-βD-βE-βF-βG (**Fig. 3E**). However, close comparison of the solution and crystal structures revealed a subtle difference, i.e., GNNV-P in solution exhibits a long flexible F’-G’ loop for aa 311-330 (**Fig. 3E**), but this loop is interrupted by a short β-strand (aa 323-326) in the crystal structure (**Fig. 3E**). Interestingly, this loop lies in the trimeric interface of the crystal structure (**Fig. S11**).

### MD simulations predict that GNNV-P forms a trimer at pH 5.0 with critical subunit interactions perturbed by site-directed mutagenesis

Although our AUC experiments clearly showed that GNNV-P molecules adopt a trimeric form at pH 5.0 (**Fig. 3A** & **Fig. S9**), the subunit interactions governing trimer formation remained unknown. Since NMR is not readily amenable to elucidating subunit interactions within a homo-oligomer, we employed an *in silico* approach using molecular dynamics (MD) simulations, representing a well-established method for understanding protein dynamics, to track GNNV-P structural transitions from pH 7.0 to 5.0. To perform this experiment, we utilized our GNNV-P solution structure determined at pH 7.0 and adjusted the protonation state of amino acid side-chain groups to pH 5.0. Three separate GNNV-P molecules without any symmetry imposed on their spatial arrangement were used as an initial point for our MD simulations, which resulted in a stable trimer (**Fig. 4A** & **B**), as evidenced by the overall Root Mean Square Deviation (RMSD) quickly reaching a stationary phase along the MD time trajectory **(Fig. S12)**. In this complex, the three GNNV-P molecules tightly associated with one another via identical interfaces between neighboring A/C, A/B and B/C subunits, with the total buried interfacial area estimated by PDBePISA as 1560 Å^2^ [30]. Moreover, three-fold rotational symmetry emerged for the arrangement of the three GNNV-P molecules, largely matching the structural arrangement of GNNV molecules in the crystal model (**Fig. S13**) [16]. In addition to predicting trimer formation, our MD simulations revealed specific interactions at the inter-GNNV-P level that could be readily verified against the NMR experimental data or tested using mutagenesis approaches.

**Figure 4.**
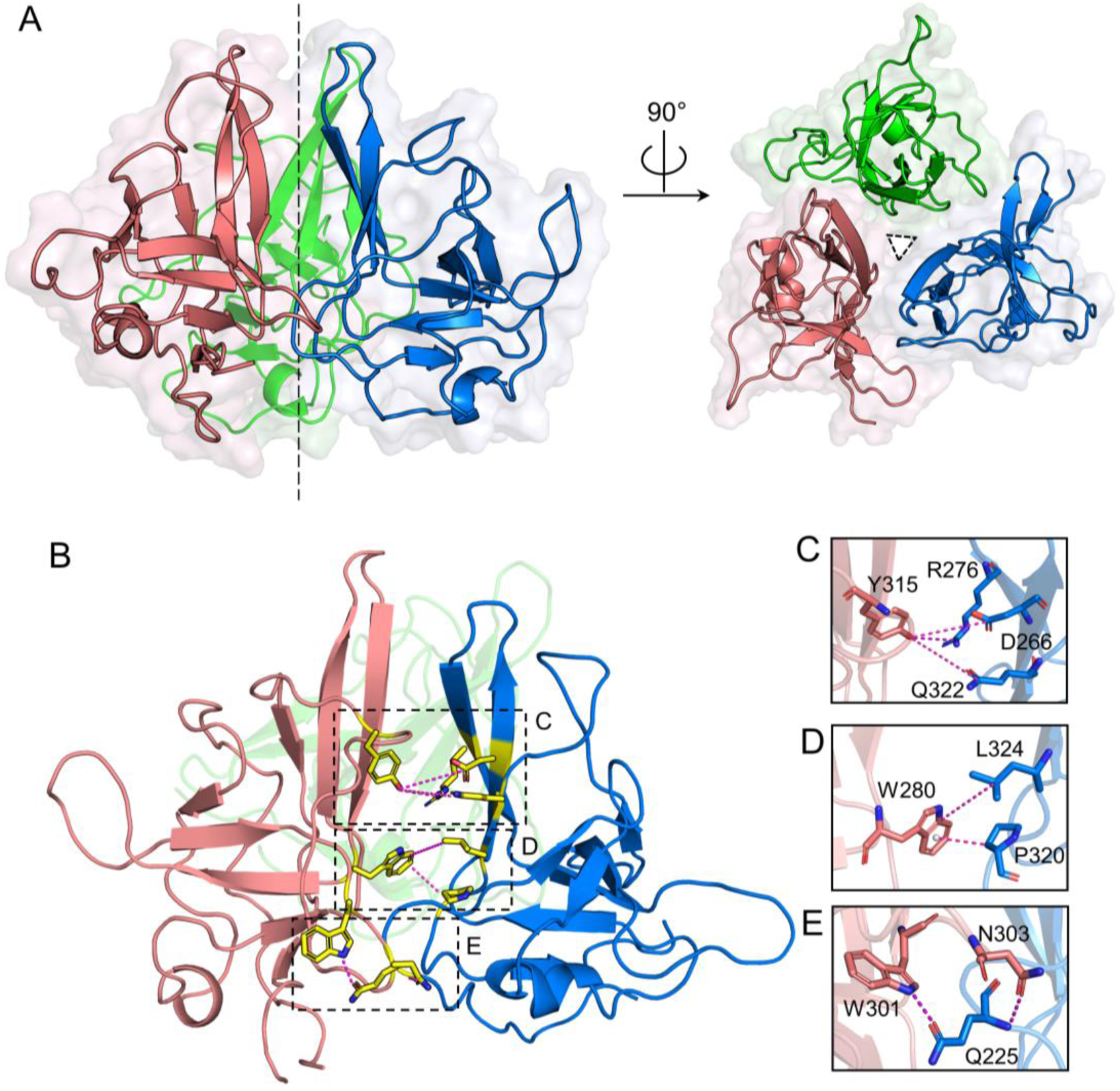
MD-predicted structure of GNNV-P oligomer at low pH. **(A)** Top and side view cartoon representations of the GNNV-P trimer formed under acidic pH conditions, as determined by molecular dynamics (MD) simulations. Chain A (blue), chain B (pink), and chain C (green). **(B)** Intermolecular interactions between neighboring GNNV-P units in the trimer. Residues at the trimeric interface are depicted as yellow sticks, and interactions are shown as dashed magenta lines. **(C-E)** Details of trimeric interface interactions between GNNV-P chains A and B, as boxed in (B).

Therefore, we searched the MD-predicted structure for residues in the trimer interface, guided by the pH-sensitive regions we had already identified (**Fig. 2C** & **D**). We detected three pairs of contacts between the side-chains of one subunit and those of a neighboring subunit: Y315 with D266/R276/Q322 (**Fig. 4C**); Q225 with W301/N303 (**Fig. 4D**); and W280 with L324/P326 (**Fig. 4E**). Notably, some of these contacts are disfavored at neutral pH, as they are sterically hindered by the F’-G’ loop. Among the three interacting pairs, the W280 with L324/P326 pair seems to be relatively strong as the interactions are dominated by hydrophobic and CH-π interactions, whereas the other two pairs represent either polar interactions or primarily hydrogen bonds. To establish if these three MD-predicted inter-GNNV-P interactions are plausible, we generated three sets of single alanine mutants and assayed their impacts on GNNV-P oligomerization by means of AUC. The first set of mutations encompassed L324A, P326A, W280A and I323A. As shown in **Fig. S14**, the L324A, P326A or W280A mutations abolished oligomer formation at low pH (5.0), indicating that this MD-predicted pair of interactions is indeed critical to stabilizing the trimer. As a control, we also mutated I323, a residue in the same pH-sensitive region, but that does not exhibit inter-molecular interactions in the MD-predicted structure nor exhibited significant CSPs. As expected, the I323A mutation did not interfere with GNNV-P oligomer formation at low pH (5.0). AUC data for the second set of mutations showed that Q322A impaired oligomerization at pH 5.0, whereas R276A still permitted trimer formation, implying that residue Q322 is critical whereas R276 is not. Regrettably, we were not able to perform AUC experiments for the third set of mutations as the W301A mutant protein failed to fold correctly. Nevertheless, our mutagenesis study verified the importance of the inter-GNNV-P interactions predicted by MD simulations, validating the MD simulations results. We also noticed a conserved histidine at position 281 of GNNV-P (**Fig. S15**), and postulated that it might serve as a histidine switch through pH-induced protonation/de-protonation, as utilized for enveloped virus activation under acidic conditions [31]. However, AUC of a H281Y mutant ruled out that this histidine is involved in low-pH-induced GNNV-P oligomerization (**Fig. S14**).

### NMR signatures of GNNV-P conformational change at low pH

Apart from capturing GNNV-P trimerization under the acidic condition, our MD simulations indicated a change in GNNV-P secondary structure at pH 5.0. Compared to backbone CSPs, C_α_ and C_β_ CSPs are more sensitive to secondary structure change. In searching for residues with pronounced C_α_ and C_β_ CSPs induced by low pH (**Fig. 5A**), we identified a sub-region within Region III (aa 312-327) (**Fig. 5A**) exhibiting an increased β-strand propensity at pH 5.0. This sub-region is composed of residues P320-L324 and it coincides precisely with an MD-predicted region that became a β-strand at pH 5.0. This region in the MD-predicted structure formed intra-molecular hydrogen bonds with residues S264 and D266 (**Fig. 5B**), two residues that belong to β-strand C (aa 263-267) (**Fig. 3E**). The impact of these contacts is reflected by the pronounced CSPs of Region I (aa 263-279) (**Fig. 3A**). Thus, details from our MD simulations could be further validated by CSP data extracted from NMR experiments. Of note, the β-strand of P320-L324 is stabilized by its association with β-strand C; these two strands together with β-strand D form a three-stranded antiparallel β-sheet in the GNNV-P structure at pH 5.0 obtained by MD simulations.

**Figure 5.**
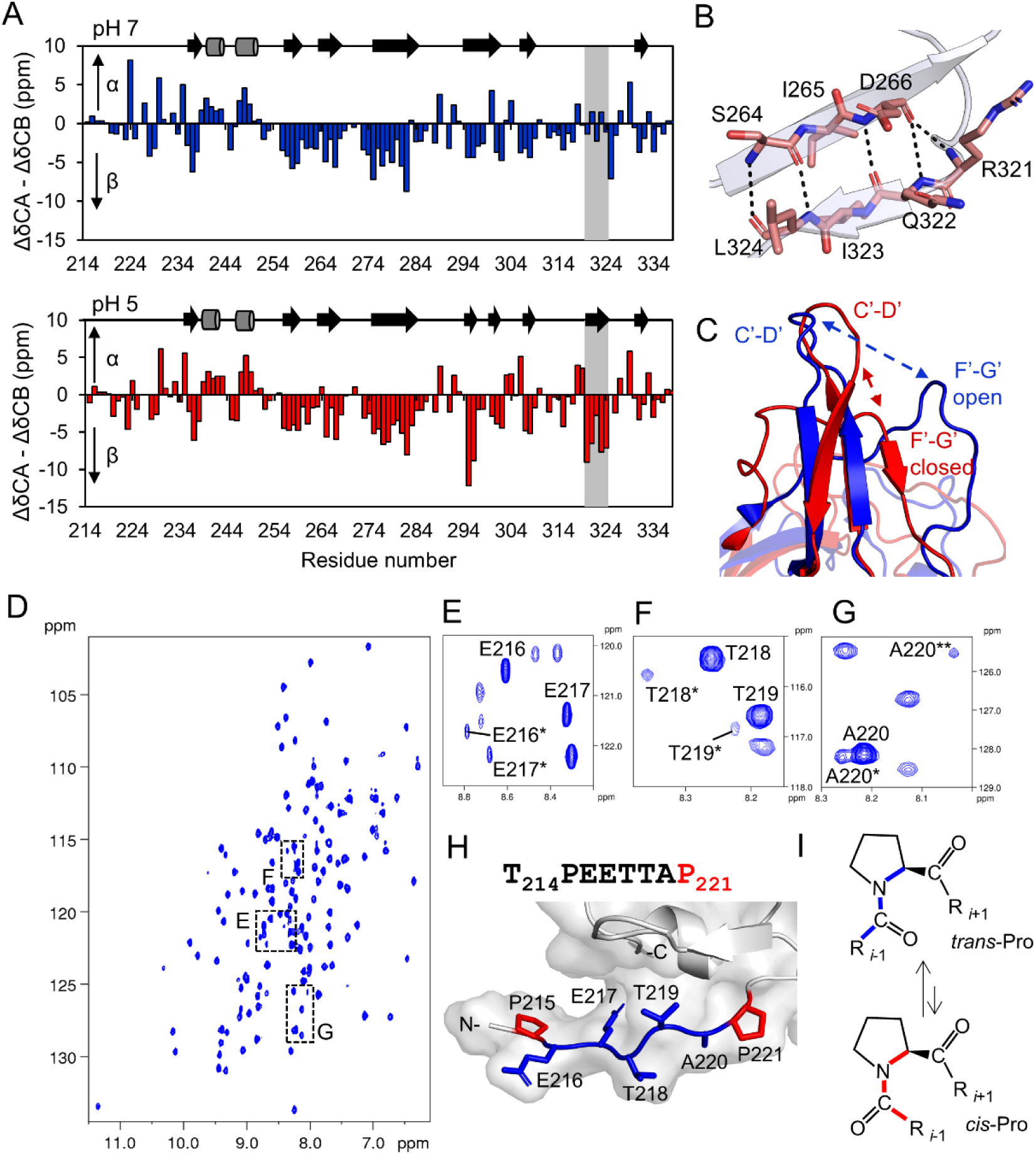
GNNV-P undergoes a low pH-induced conformational change. **(A)** Secondary structure propensities of GNNV-P at pH 7.0 and pH 5.0, calculated using secondary chemical ΔC_α_ – ΔC_β_ shifts and plotted against amino acid sequence. The Q320-P324 region showing an increased β-strand propensity at pH 5.0 is highlighted in grey. The secondary structures are shown above each chart, with arrows and cylinders representing β-strands and α-helices, respectively. **(B)** MD simulations revealed formation of a short β-strand (comprising residues Q322-L324) within the F’-G’ loop at low pH. Hydrogen bonds between R321-L324 and D266-S264 were identified as stabilizing this region. **(C)** Superimposition of GNNV-P at pH 7.0 (NMR structure, blue) and pH 5.0 (MD model, red) showing the open and closed conformations of the F’-G’ loop, respectively. **(D)** ^1^H-^15^N HSQC spectra of U-D-, ^13^C-, and ^15^N-labeled GNNV-P at pH 5.0. The regions with duplicate signals for residues 216-220 in the linker region are marked with black boxes. **(E-G)** Details of the U-D-, ^13^C-, and ^15^N-labeled GNNV-P ^1^H-^15^N HSQC spectra highlighted by black boxes in D. **(H)** Sequence of the N-terminal linker T214-P221 and a stick representation of the linker region with proline residues colored red and residues presenting duplicate NMR signals colored blue. **(I)** Schematic representation of *cis*-*trans* isomerization of an Xaa-Pro peptide bond. Proline residues can switch between the *trans* (blue) and *cis* (red) conformations.

Given the aforementioned conformational change in the F’-G’ loop at low pH (**Fig. 3E**), the space between the F’-G’ and the C’-D’ loops collapsed (**Fig. 5C**). Consequently, a pocket in GNNV-P open at neutral pH becomes closed at acidic pH, rendering individual GNNV-P units more compact, as evidenced by a decrease in surface area from 14053 Å^2^ (pH 7.0) to 12160 Å^2^ (pH 5.0) calculated using Pymol [32]. Since the β-strand of P320-L324 is situated in the center of the trimer (**Fig. S12)**, its switching from the loop configuration mitigates inter-GNNV-P steric hindrances that, together with neutralization of interfacial repulsing electrostatic potential at pH 5.0 (**Fig. S16**), results in a compact trimer. This detailed mechanism nicely explains the low-pH-induced compaction of protrusions on GNNV particles observed by cryo-EM (**Fig. 1A & B**).

To understand the linker malleability proposed by our cryo-EM observations, we searched the NMR data for residues in the linker region (214-220 aa). During our assignments of NMR spectra for GNNV-P at pH 5.0, we observed duplicate NMR signals (**Fig. 5D-G**) for residues preceding the proline residue (P221) situated at the junction between the linker and P-domain. This result indicates that the linker gradually shifts among different conformations, possibly due to the peptide bond of P221 undergoing *cis*-*trans* isomerization at pH 5.0 (**Fig. 5H-I**). Our MD simulations support this conformational change of the linker around P221 (**Fig. S17**). However, due to peptidyl bond isomerization being a slow process, directly capturing the switching from a *trans* to *cis* conformation was not possible with our MD simulations. Although the *trans* configuration is favored for peptide bonds, interconversion between the *cis* and *trans* configurations for a Xaa-Pro peptide bond has been previously demonstrated [33]. In addition, an increased population of P221 in *cis* conformation at low pH has also been inferred by analyzing our ^13^C chemical shifts data using the PROMEGA webserver [34]. An additional NMR signature for malleability of the linker is evidenced by the signal for residue A220 in 2D HSQC spectra at pH 5.0, which exhibited various C_α_ and C_β_ chemical shifts in the HNCACB spectra (**Fig. S18**). Taken together, our NMR results provide spectral evidence to support that the linker between the P-domain and S-domain of GNNV is malleable.

### Interactions of GNNV-P with host surface glycan receptors

Upon encountering a host cell, the protrusions of GNNV-P represent an immediate viral structural motif for the virus to engage with receptors on the host cell surface. Cell surface glycans such as sialic acids have been reported as common cell receptors for distinct NNVs [35]. By using molecular docking analysis, Nishizawa et al. identified interactions between the terminal moiety of sialic acid and a GNNV protrusion [36]. That analysis was based on the then available three-dimensional structure of the GNNV P-domain as a compact trimer obtained from X-ray crystallography [16]. However, our NMR solution structure indicates that the GNNV P-domain adopts a monomeric form in the pH conditions when GNNV engages with host cell surfaces.

Thus, using our P-domain structure at pH 7.0, we could re-evaluate potential modes of interaction between sialic acids and GNNV-P. To do so, we conducted a molecular docking analysis with our solution structure of GNNV-P (pH 7.0) against two sialoside isomers, i.e., Neu5Ac-(α2,3)-Lac and Neu5Ac-(α2,6)-Lac. Remarkably, as shown in **Fig. 6A**, the terminal Neu5Ac of both ligands could be inserted deep into the pocket formed by the F’-G’ loop (**Fig. 5C**). The *N*-acetyl chain of Neu5AC was located inside the binding pocket and the *N*-acetyl methyl group was oriented towards the hydrophobic part of this pocket (**Fig. 6B & C**), with Neu5Ac binding being further stabilized by a number of interactions with the conserved residues lining the pocket. Those contacts include hydrogen bonds between R261 and the sialic acid carboxylate and hydroxyl groups of the penultimate galactose. The N-acetyl group of Neu5Ac-(α2,3)-Lac was stabilized by hydrogen bonds with S264 and Q319, and another hydrogen bond formed between the hydroxyl group of the penultimate galactose and Q322 (**Fig. 6B**). For Neu5Ac-(α2,6)-Lac, sialic acid formed additional interactions with S264, Q322, and I323 (**Fig. 6C**).

**Figure 6.**
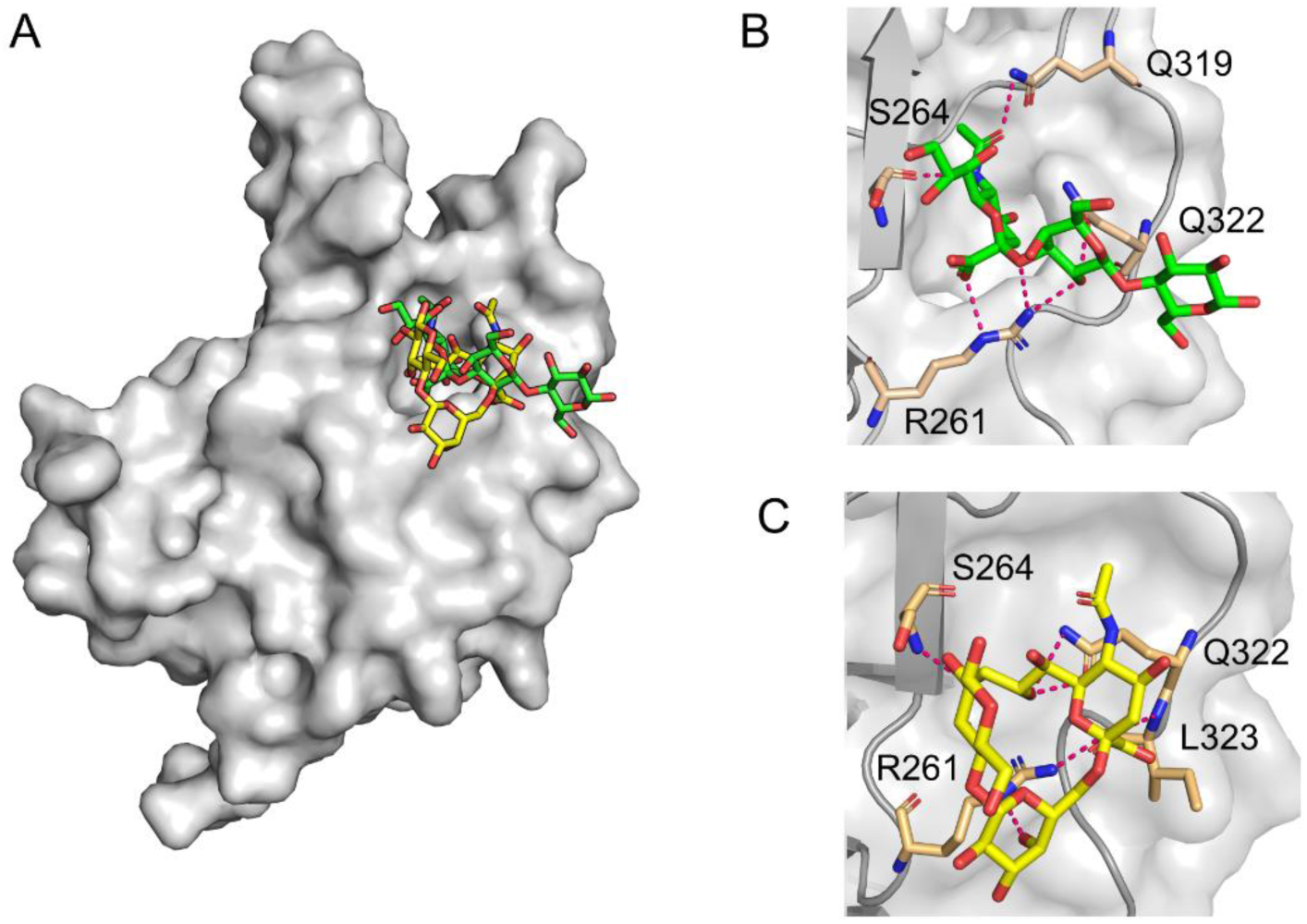
Model of Neu5Ac-Lac binding to GNNV-P at neutral pH. **(A)** Surface representation of GNNV-P monomer in complex with Neu5Ac-(α2,3)-Lac (green) and Neu5Ac-(α2,3)-Lac (yellow). **(B)** Detail of Neu5Ac-(α2,3)-Lac binding to GNNV-P in the open pocket conformation. **(C)** Detail of Neu5Ac-(α2,6)-Lac binding to GNNV-P in the open pocket conformation. In B and C, residues interacting with Neu5Ac-Lac are shown as sticks, and selected contacts between GNNV-P and Neu5Ac-Lac are represented by pink dashed lines.

Together, these specific interactions endow a strong binding affinity between GNNV-P and Neu5Ac-(α2,3)-Lac or Neu5Ac-(α2,6)-Lac, with free energies of -6.42 kcal/mol and -6.48 kcal/mol, respectively. Significantly, those interactions could be augmented multivalently by the three GNNV P-domains that form a viral capsid protrusion. As a control, we also performed molecular docking analysis using our trimeric structure (pH 5.0) against the same two sialoside isomers. Since the pockets in the trimeric structure are closed at this pH due to a conformational switching in the F’-G’ loop, Neu5Ac-(α2,3)-Lac or Neu5Ac-(α2,6)-Lac could no longer bind in the pocket. Instead they attached to the pocket vestibule at the apex of the GNNV-P trimer (**Fig. S19**), confirming the docking results of Nishizawa et al. [36].

## Discussion

Attachment of a virion to a host cell surface represents the first and a critical step during virus infection. Compared to other viruses, GNNVs dock on their host cells, e.g., Grouper fin cells, under slightly alkaline conditions because groupers live in relatively warm oceanic regions that exhibit alkaline pH of 8.1. Thus, a GNNV that has entered a host cell and is trafficked into an endosome (pH 5.5-5.0) encounters a wide range of pH variations. How the structures of GNNVs respond to such drastic pH changes during host cell entry and genome release had been virtually unknown, with the structures assumed to be akin to those revealed by the previously generated crystal structure of GNNV VLP [16]. Herein, we have challenged that notion by investigating the structures of native GNNV virus and VLP in various pH conditions. To do so, we deployed cryo-EM, an appropriate method for high-resolution structural determinations of non-crystal specimens and adaptable to a wide range of aqueous conditions. Surprisingly, we observed that the protrusions on the GNNV capsid could switch configuration in a pH-dependent manner, adopting an erect position at neutral and alkaline pH, but resting on the capsid shell at acidic pH (5.0) with concurrent compaction and rotation (**Fig. 1B**). We next expressed the P-domain alone and used analytical ultracentrifugation to uncover that this GNNV-P protein is largely monomeric at neutral pH, but assembles into a trimer at acidic pH. Further analysis of P-domain pH sensitivity using solution NMR combined with MD simulations allowed us to resolve the structure of GNNV-P in solution, which revealed an unusual pH-dependent mechanism of conformational switching for a long flexible loop comprised of 20 amino acids (aa 311-330). This F’-G’ loop forms a loop at pH 7.0, but part of it converts to a short β-strand at pH 5.0. These two conformations play a pivotal role in controlling the opening and closing of a pocket formed between the F’-G’ and C’-D’ loops of the P-domain (**Fig. 5C**). At neutral pH, the F’-G’ and C’-D’ loops lie far apart, so the pocket is open, whereas it becomes closed in acidic pH because part of the F’-G’ loop converts to a β-strand, forming a three-stranded antiparallel β-sheet with other β-strands. Concurrently, this conformational switching ameliorates loop-elicited steric hindrance between neighboring P-domains that prevent P-domain trimerization (**Fig. 4**). The P-domain trimer is further stabilized by inter-GNNV-P interactions at their interface. Those interactions encompass critical contacts between residues P326 and W280 of neighboring GNNV-P, as verified by site-directed mutagenesis. In contrast to the crystal structure [16], the GNNV-P trimer in solution is stabilized solely by protein-protein interactions since the solution does not contain any divalent ions used for crystallization [16]. Moreover, unlike the GNNV-P structure determined by X-ray crystallography, our NMR analyses allowed us to address the molecular behavior of the linker connecting the P-domain to the S-domain, which revealed its pH-dependent malleability that underlies motion of the protrusions. The NMR spectra of this linker revealed that it can adopt multiple conformations in acidic conditions via peptide bond isomerization of a conserved proline (P221) at the junction of the linker and P-domain. Taken together, our NMR and MD analyses establish the pH-dependent movement, compaction, and trimerization of P-domains that explains the conformational change revealed by cryo-EM of GNNV particles.

Given that our NMR solution analysis revealed the structure of GNNV-P under conditions compatible with virus entry, the resulting solution structure (monomer) represents a more appropriate system (relative to the trimeric crystal structure) for evaluating how NNVs interact with host receptors. Our molecular docking analysis on GNNV-P monomer interacting with a sialic acid molecule demonstrated that the open pocket formed by the F’-G’ loop interacts with host surface glycans. As the tested sialic acids form contacts with highly conserved residues deep in the pocket, our results consolidate the notion that sialic acids are utilized as common cell receptors across all NNV genotypes [35]. Since a common cell receptor cannot differentiate among NNVs, a protein co-receptor is required to determine host specificity [37–39]. Such protein co-receptors include 90ab1 [38] and nectin [39], which bind to amino acids 213-230 and 221-238, respectively, of the P-domain. Interestingly, both these host-determining regions of NNVs encompass or overlap with a highly variable sequence (aa 223-244) [40]. As this variable sequence lies close to the linker (aa 214-220), yet distant from the sialic acid-binding pocket, we speculate that the pH-dependent conformational change of the linker revealed by our structural findings plays a role in dynamically controlling the accessibility of those host-determining regions. Based on the interactions between the protrusion pocket and sialic acid receptors, as well as those between the protrusion host-determining region and protein co-receptors, we propose a pH-dependent NNV-host engagement model (**Fig. 7**) relevant to virus entry and perhaps subsequent host receptor escape [41]. These novel insights into GNNV structural dynamics should prove useful in facilitating the design of vaccines against NNVs [17, 42–44].

**Figure 7.**
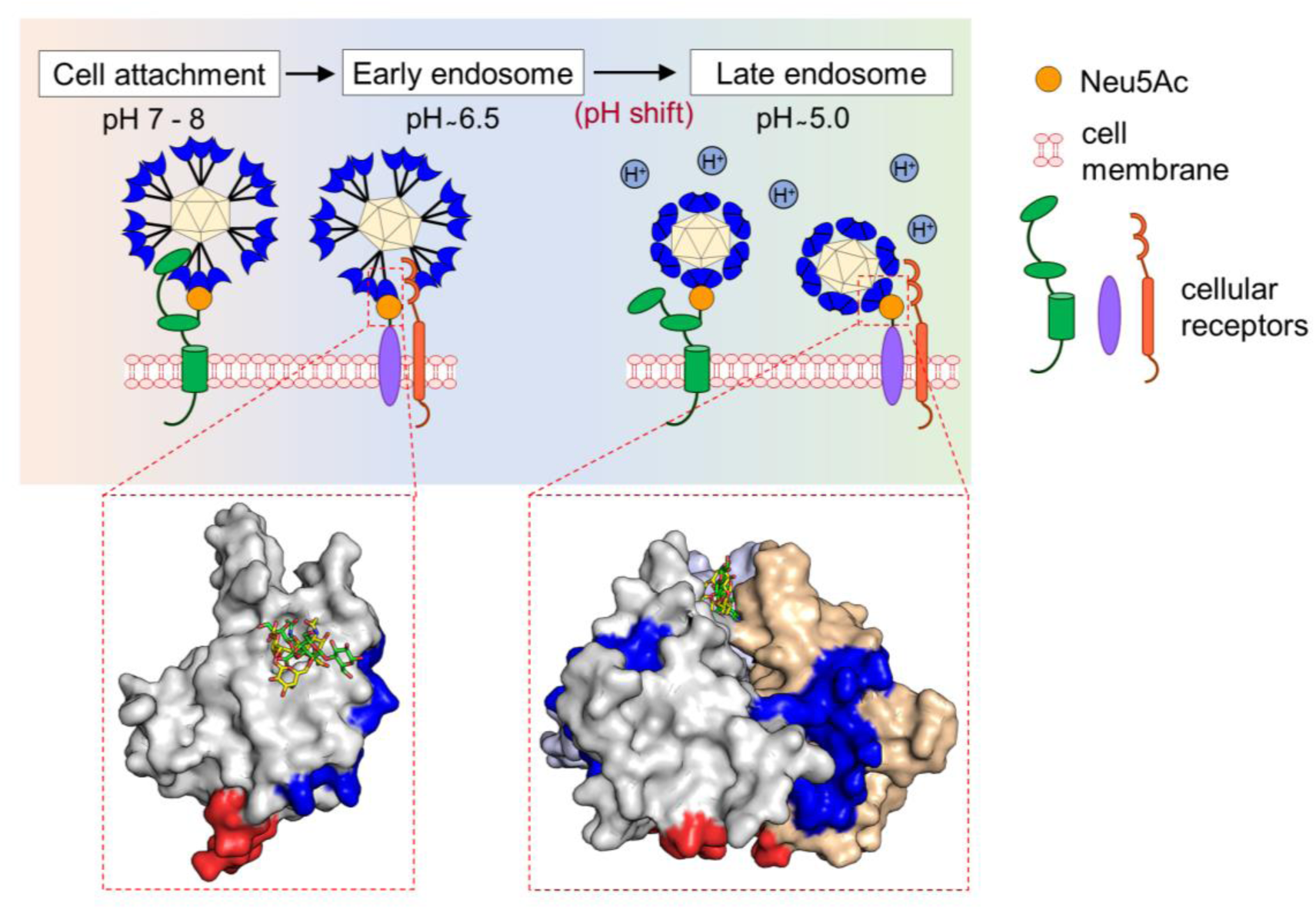
Model of GNNV interactions with host cell surface receptors. During infection, GNNV attaches to cells by interacting with sialic acid moiety on the host cell surface and with cellular receptors (HSP70, HSP90ab1, or nectin-4). At alkaline or neutral pH, protrusions on the GNNV surface comprising P-domains are in extended and loose configuration, with the sialic acid binding pocket being open, and linker region (red) and host-determining region (blue) accessible for interactions with cellular receptors. Acidification in late endosomes induces conformational change of GNNV P-domain to result in more compact structure, promoting the formation of a compact trimer of P-domains. At the resting/prone position, protrusions on the GNNV surface are with sialic acid binding pocket being closed that alternative sialic acid binding site is rearranged to the tip of the protrusions, and they conceal the linker region of GNNV P-domain, leading to potential detachment of those cellular receptors.

Like GNNV, norovirus is a non-enveloped virus with pronounced surface protrusions. Each norovirus protrusion is composed of two copies of the P-domain, which is connected to an S-domain via a flexible hinge [45]. Recently, Song et al. investigated mouse norovirus by cryo-EM under various pH conditions [46], and revealed that mouse norovirus protrusions rest on the virus capsid in neutral pH (< 8.0), but become erect at alkaline pH (> 8.0). This dynamic switching of protrusion configurations apparently governs viral infection, as the resting protrusions appear to be more accessible to cellular receptors [46]. Our cryo-EM structures of GNNV demonstrate that its capsid protrusions behave similarly to those of norovirus in terms of changing in response to altered pH. However, the pH thresholds for the conformational change differ significantly. In addition, the infectious form of GNNV may differ from that of norovirus with respect to the protrusion positions. For GNNV, the infectious form is more likely to be represented by erect protrusions, unlike the prone protrusions utilized by norovirus. The ability of protrusions to switch positions shared by these two viruses is likely attributable to a common feature of linker malleability. For GNNV, and probably NNV in general, this malleability is endowed by a conserved proline (P221), the peptide bond of which can isomerize between *cis* and *trans* configurations (**Fig. 5**). Noroviruses host a PPT (Pro-Pro-Thr) sequence in its hinge region that is pivotal for protrusion malleability [45], but the behavior of the PPT sequence in response to changing pH awaits investigation.

In summary, our analyses of GNNV using cryo-EM and NMR, combined with MD simulations and mutagenesis, have uncovered a unique pH-sensing mechanism underlying the dynamics of its capsid protrusions. There are three novel and significant outcomes from our study. First, we show that the protrusions on native GNNV virions can switch from an erect to a prone configuration at low pH, further advancing from the static VLP crystal structure [16]. Second, we have determined by NMR the structure of the GNNV P-domain in solution at pH 7.0. Third, the morphological change of the capsid protrusions is attributable to a pH-dependent conformational change in a loop of the GNNV P-domain, with a malleable linker being responsible for the protrusion movement. In conclusion, our work on GNNV protrusions has unveiled a structural mechanism utilized by a betanodavirus governing its infectivity, which is similar to that deployed by a phylogenetically distant non-enveloped human virus, implying that the similarity in linker malleability might have been acquired through convergent evolution.

## Materials and Methods

### Sample preparation of GNNV and VLPs for cryo-EM

GNNV and VLPs were purified using a 10-40% (w/w) sucrose density gradient, as previously described in [8] and [25] respectively. To prepare GNNV and VLPs in different pH conditions, the purified particles were pelleted down by ultra-centrifugation at 30,000 rpm for 3.5 hours at 4 °C (Beckman Coulter, OptimaTM L-90K Ultracentrifuge, rotor: SW 41 Ti). The particle pellets were then re-suspended overnight in 100 μl of TN buffer (50 mM NaCl, 50 mM Tris-HCl, pH 8.0), MES buffer (50 mM NaCl, 50 mM MES, pH 6.5), or acetate buffer (50 mM NaCl, 50 mM sodium acetate, pH 5.0), respectively.

To prepare the cryo-EM samples of GNNV and VLPs, approximately 3.5 μl of protein solution was deposited onto a Quantifoil R1.2/1.3 holey carbon grid (Quantifoil Micro Tools GmbH, Jena, Germany) coated with a thin carbon film. The grid was then rapidly plunged into liquid nitrogen-cooled liquid ethane and stored in liquid nitrogen until imaging. The cryo-EM grids of GNNV at pH 6.5 and pH 5.0 were prepared using a Vitrobot Mark IV system (Thermo Fisher Scientific, Hillsboro, OR, USA) at 4 °C and 100% humidity, with a blotting time of 3.5 seconds. For VLPs at pH 8.0, pH 6.5 and pH 5.0, the cryo-EM grids were prepared using a Leica EM GP system (Leica Biosystems, Deer Park, IL, USA) with the sensor off option—the cryo-EM grids of VLPs at pH 8.0 and pH 5.0 were prepared at 15 °C and 80% humidity, with a blotting time of 1.2 seconds; whereas the cryo-EM grids of VLPs at pH 6.5 was prepared at 22 °C and 95% humidity, with a blotting time of 0.5 seconds. All subsequent steps were conducted at liquid nitrogen temperature to prevent de-vitrification.

### Cryo-EM data acquisition

The cryo-EM grids containing GNNV virions at pH 6.5 and pH 5.0 were examined using cryo-ARM (JEOL Ltd., Akishima, Tokyo, Japan) at magnifications of 40,000x and 50,000x, respectively, with pixel sizes of 1.36 Å/pixel and 1.09 Å/pixel. Cryo-EM images of GNNV at pH 6.5 and pH 5.0 were recorded using a K2 camera (Gatan Inc., Pleasanton, CA, USA) in counting mode, with an exposure time of 8 seconds for 40 frames. The dose rate was approximately 6.5 electrons/Å^2^ per second, resulting in a total accumulated dose of around 52 electrons/Å^2^ (equivalent to approximately 1.3 electrons/Å^2^ per frame).

The cryo-EM grids containing VLPs at pH 6.5 were examined using Technai F20 (FEI, Hillsboro, OR, USA) at a nominal magnification of 29,000x, resulting in a pixel size of 1.24 Å/pixel. Cryo-EM images of the VLPs at pH 6.5 were recorded using a K2 camera (Gatan Inc., Pleasanton, CA, USA) in counting mode, with an exposure time of 10 seconds for 50 frames. The dose rate was approximately 5 electrons/Å^2^ per second, resulting in a total accumulated dose of around 50 electrons/Å^2^ (equivalent to approximately 1.0 electron/Å^2^ per frame).

The cryo-EM grids containing VLPs at pH 8.0 and pH 5.0 were examined using JEM-2100F with a high-contrast pole piece (JEOL Ltd., Akishima, Tokyo, Japan) at a magnification of 50,000x, with a pixel size of 1.16 Å/pixel. Cryo-EM images of the VLPs at pH 8.0 and pH 5.0 were recorded using a DE-20 camera (Direct Electron LP, San Diego, CA, USA) in linear mode, with an exposure time of 1.5 seconds for 38 frames. The dose rate was approximately 20 electrons/Å^2^ per second, resulting in a total accumulated dose of around 30 electrons/Å^2^ (equivalent to approximately 0.8 electrons/Å^2^ per frame). The parameters for cryo-EM data acquisition are summarized in **Table 1**.

**Table 1.** GNNV cryo-EM data collection, refinement and validation statistics.

|  | **GNNV**<br>**Virion**<br>**(pH6.5)**<br>**(EMD-39215)**<br>**(PDB-8YF9)** | **GNNV**<br>**Virion**<br>**(pH5.0)**<br>**(EMD-39217)**<br>**N/A** | **GNNV**<br>**VLP**<br>**(pH8.0)**<br>**(EMD-39212)**<br>**(PDB-8YF6)** | **GNNV**<br>**VLP**<br>**(pH6.5)**<br>**(EMD-39213)**<br>**(PDB-8YF7)** | **GNNV**<br>**VLP**<br>**(pH5.0)**<br>**(EMD-39214)**<br>**(PDB-8YF8)** |
| --- | --- | --- | --- | --- | --- |
| **Data collection** |  |  |  |  |  |
| EM equipment | Cryo-ARM | Cryo-ARM | JEM-2100F | FEI F-20 | JEM-2100F |
| Voltage (kV) | 200 | 200 | 200 | 200 | 200 |
| Cs (mm) | 1.4 | 1.4 | 3.3 | 2.3 | 3.3 |
| Magnification (nominal) | 40,000 | 50,000 | 50,000 | 29,000 | 50,000 |
| Detector | K2 | K2 | DE-20 | K2 | DE-20 |
| Detector (Operation mode) | Counting | Counting | Linear | Counting | Linear |
| Dose rate (e<sup>-</sup>/Å<sup>2</sup> per second) | ~ 6.5 | ~ 6.5 | ~ 20 | ~ 5 | ~ 20 |
| Pixel size (Å) | 1.36 | 1.09 | 1.16 | 1.24 | 1.16 |
| Electron exposure (e<sup>-</sup>/Å<sup>2</sup>) | ~ 52 | ~ 52 | ~ 30 | ~ 50 | ~ 30 |
| Exposure time (s) | 8 | 8 | 1.5 | 10 | 1.5 |
| Frames (no.) | 40 | 40 | 38 | 50 | 38 |
| Defocus range (μm) | -0.45 ~ -3.05 | -0.42 ~ -4.18 | -0.57 ~ -2.76 | -0.40 ~ -2.91 | -0.87 ~ -2.93 |
| **Reconstruction** |  |  |  |  |  |
| Software | CryoSPARC<br>(v2.0) | CryoSPARC<br>(v2.0) | CryoSPARC<br>(v4.3) | CryoSPARC<br>(v4.3) | CryoSPARC<br>(v4.3) |
| Micrographs stacks (no.) | 391 | 577 | 588 | 1,009 | 265 |
| Final particle images (no.) | 9,626 | 26,494 | 40,334 | 39,884 | 22,346 |
| Symmetry imposed | I | I | I | I | I |
| Map final resolution (Å) † | 3.12 | 4.36 | 3.23 | 2.82 | 3.52 |
| Map sharpening B-factor (Å<sup>2</sup>) | -82.6 | -197.8 | -154.7 | -110.5 | -158.5 |
| **Atomic modeling** |  |  |  |  |  |
| Software | Coot &<br>Phenix | - | Coot &<br>Phenix | Coot &<br>Phenix | Coot &<br>Phenix |
| Number of protein residues <sup>#</sup> | 509 | - | 509 | 509 | 873 |
| Number of metal ions <sup>#</sup> | 3 (Ca<sup>2+</sup>) | - | 3 (Ca<sup>2+</sup>) | 3 (Ca<sup>2+</sup>) | 3 (Ca<sup>2+</sup>) |
| Number of atoms <sup>#</sup> | 3,895 | - | 3,895 | 3,895 | 6,748 |
| Map CC (around atoms) <sup>#</sup> | 0.83 | - | 0.89 | 0.87 | 0.82 |
| RMSD bond lengths (Å) <sup>#</sup> | 0.004 | - | 0.005 | 0.005 | 0.005 |
| RMSD bond angles (°) <sup>#</sup> | 1.009 | - | 0.992 | 0.650 | 0.985 |
| Clash score <sup>#</sup> | 8.76 | - | 7.09 | 9.67 | 14.48 |
| Ramachandran favored (%) <sup>#</sup> | 96.22 | - | 96.62 | 97.42 | 96.08 |
| Ramachandran allowed (%) <sup>#</sup> | 3.78 | - | 3.38 | 2.58 | 3.92 |
| Ramachandran outliers (%) <sup>#</sup> | 0 | - | 0 | 0 | 0 |
| Rotamer outliers (%) <sup>#</sup> | 0 | - | 0 | 0 | 0 |
| C<sub>β</sub> deviations <sup>#</sup> | 0 | - | 0 | 0 | 0 |
| MolProbity score <sup>#</sup> | 1.73 | - | 1.61 | 1.62 | 1.94 |

<sup>#</sup> Statistics are given for one icosahedral asymmetric unit
<sup>†</sup>According to Fourier shell correlation (FSC)=0.143.

### Single-particle image processing and 3D reconstruction

All cryo-EM image stacks underwent motion correction and dose weighting using MotionCor2 [47] with a 5 x 5 patch. The contrast transfer function (CTF) was determined using CTFFIND4 [48] from the motion-corrected and dose-weighted images. Particle picking was conducted in cryoSPARC [49] using 2D templates generated from a previously determined VLP cryo-EM map [25]. After particle extraction and removal of bad particles through 2D classification, the remaining particles were used for further *ab initio* reconstruction and heterogeneous refinement with icosahedral symmetry (I). A subset of particles from a good 3D class with more particles and better resolution were chosen for homogeneous refinement with icosahedral symmetry (I). Overall resolution was assessed using the Fourier Shell Correlation (FSC) = 0.143 criterion, and local resolution was calculated within cryoSPARC [49]. The final cryo-EM maps achieved overall resolutions of 3.12 Å (GNNV pH 6.5), 4.36 Å (GNNV pH 5.0), 3.23 Å (VLP pH 8.0), 2.82 Å (GNNV pH 6.5), and 3.52 Å (GNNV pH 5.0), respectively (**Fig. S3** & **S4, and Table 1**). Visualization of the resulting 3D density maps was performed using UCSF Chimera [50]. The details of single-particle image reconstructions of GNNV and VLPs can be found in the flowcharts in **Fig. S20** & **S21**, and the cryo-EM reconstruction details are summarized in **Fig. S3** & **S4**. Additional information regarding cryo-EM reconstruction statistics is available in **Table 1**.

### Protein construct and site-directed mutagenesis

The cDNA encoding the Dragon grouper NNV P-domain (aa 214-338, GNNV-P) was tagged with His_6_-yeast SUMO (Smt3) at the N-terminus and was cloned into pETDuet-1 vector. GNNV-P mutant constructs—namely R276A, W280A, H281Y, W301A, Q322A, I323A, L324A, and L325A—were created using plasmids harboring the wild type GNNV-P sequence and a QuickChange Lightning site-directed mutagenesis kit (Agilent Technologies, CA, USA). Mutations were confirmed by PCR sequencing (Genomics Inc., Taiwan). Primers used for mutagenesis were synthesized by Tri-I Biotech Inc. (NTC, Taiwan).

### Protein expression and purification

Wild type and mutant GNNV-P proteins were expressed using transformed *Escherichia coli* BL21 (DE3) strain according to expression and purification protocols reported previously [27]. U-[^2^H, ^13^C, ^15^N] triple-labeled proteins for NMR assignments at pH 5.0 were expressed in M9 minimal medium prepared using 100% D_2_O (Sigma-Aldrich) as solvent (M9 D_2_O medium), with 1 g/L ^15^NH_4_Cl (Sigma-Aldrich) and 2 g/L U-^13^C_6_-Glucose (Cambridge Isotope Laboratories) as the sole nitrogen and carbon sources, respectively, according to a modified expression protocol. In brief, overnight culture of transformed *E. coli* was first transferred to 1L of LB medium supplemented with 100 μg/ml ampicillin and grown until the OD_600_ reached a value of 1.0. The cells were then collected by centrifugation and excess LB medium was removed before re-suspending the cells in M9 D_2_O medium. Protein expression was induced by addition of IPTG (final concentration of 0.3 mM) dissolved in 100% D_2_O. Then, the cells were grown for an additional 4 hours at 37 °C with shaking at 150 rpm, before being harvested by centrifugation at 8000 rpm for 25 minutes at 4 °C.

### Sedimentation velocity analytical ultracentrifugation (SV AUC)

The sedimentation velocity experiments were performed in a Beckman Coulter ProteomeLab XL-I analytical ultracentrifuge equipped with a 190-800 nm absorbance optical system. The concentration of protein samples used for SV AUC was in the 0.25 - 0.4 mg/ml range. The SV experiments were carried out at a rotor speed of 60,000 rpm at 20 °C and absorbance was monitored at λ = 280 nm. Protein sample buffer was used as a reference. Sedimentation coefficients were determined using the SEDFIT v16-1c software [51]. The sedimentation velocity data was fitted into a continuous c(s) distribution model based on solving the Lamm equation by the least-squares technique [51]. Buffer density (ρ), viscosity (η), and molecule partial specific volume were estimated using SEDNTERP software [52].

### NMR spectroscopy and structure determination

All NMR experiments were performed at 298 K on Bruker AVANCE 600, 800, and 850 MHz spectrometers equipped with 5 mm triple resonance TXI cryogenic probes including a shielded Z-gradient. Samples containing 10% D_2_O were loaded into 5 mm Shigemi NMR tubes for NMR experiments.

The sequence-specific backbone resonance assignments at pH 5.0 were achieved using 1.0 mM of U-[^2^H, ^13^C, ^15^N]-labeled GNNV-P protein in 20 mM sodium acetate (pH 5.0), 50 mM NaCl, 0.5 mM ethylenediaminetetraacetic acid (EDTA), 0.02% sodium azide, and 90% H_2_O/10% D_2_O. NMR spectra were processed using Bruker Topspin 3.6 and analyzed using NMRviewJ 9.2.0. [53]. The sequence-specific backbone assignments have been determined by independent connectivity analysis of HNCACB, HNCO and HN(CA)CO experiments. We completed backbone assignments for 123 out of 125 residues (98.3%), with the exceptions of Thr214 and Leu325. These resonance assignments have been deposited into the Biological Magnetic Resonance Databank with accession code 52218.

GNNV-P structural calculations at neutral pH were carried out in XPLOR-NIH software version 3.8 [54] in NMRbox [55] using experimentally determined distance restraints, hydrogen bonds, and predicted dihedral angles. Backbone and side-chain NMR assignments of the NNV P-domain at neutral pH were reported previously [27]. NOE distance restraints were derived from an ^15^N-edited NOESY-HSQC spectrum and they were analyzed using NMRviewJ software [53]. Hydrogen bonds were identified based on hydrogen/deuterium exchange experiments. The Backbone dihedral angle restraints Φ and Ψ were predicted from chemical shifts using the TALOS+ webserver [56]. For GNNV-P structural determination, 100 structures were generated according to a standard simulated annealing protocol. Twenty structures with the lowest energy were selected for refinement using the implicit solvation potential and effective energy function (EFFx) in XPLOR-NIH [57]. The twenty structures with the lowest energy and without reported violations were selected for assessment and quality checking using the protein structure validation suite in the wwPDB Validation server [58]. The final ensemble of 20 structural conformations of GNNV-P has been deposited in the Protein Data Bank (PDB entry 8XID) and the respective structural statistics are summarized in **Table 2**.

**Table 2.**
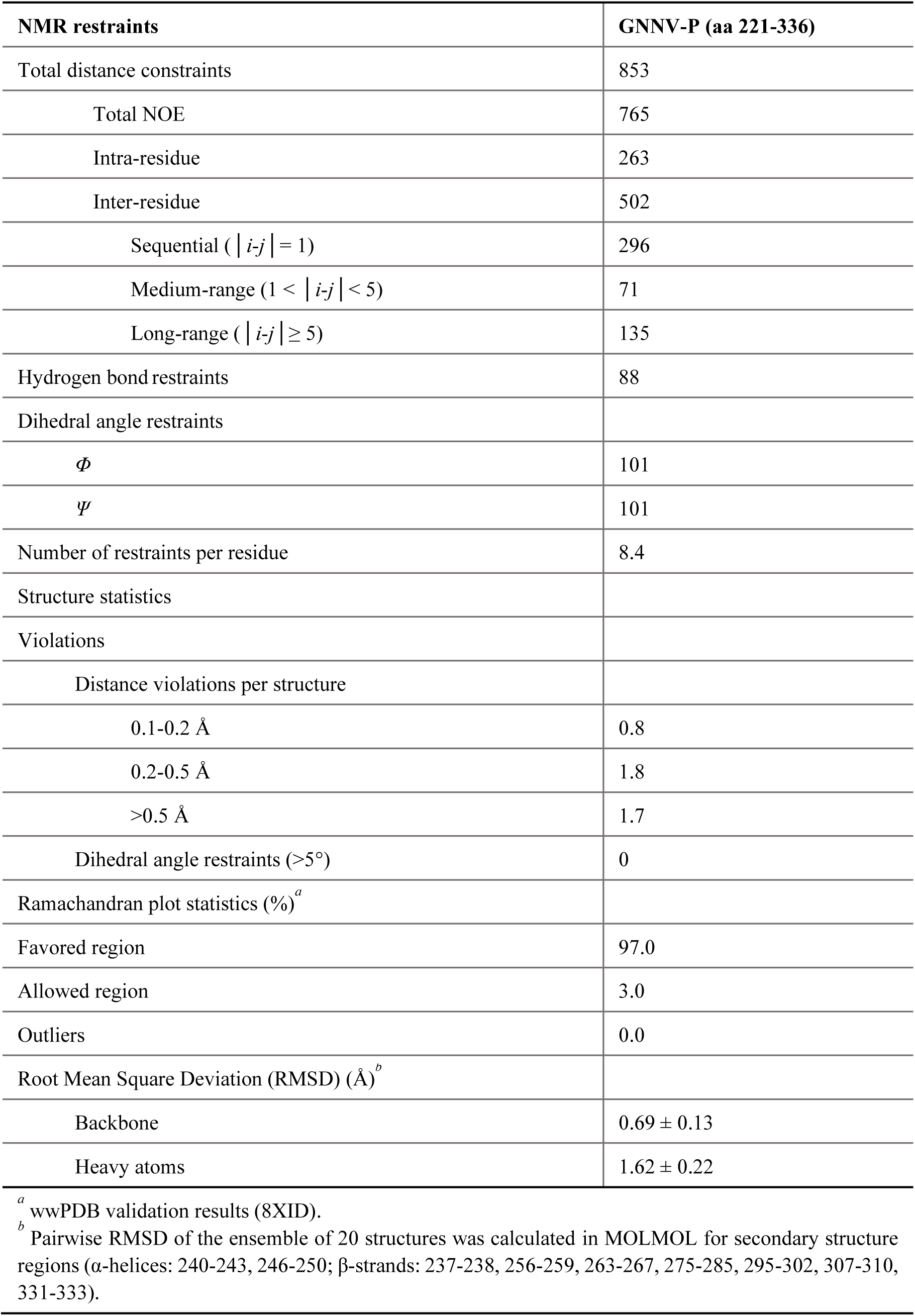
Structural statistics of the 20 lowest energy GNNV-P structures.

**Table 3.**
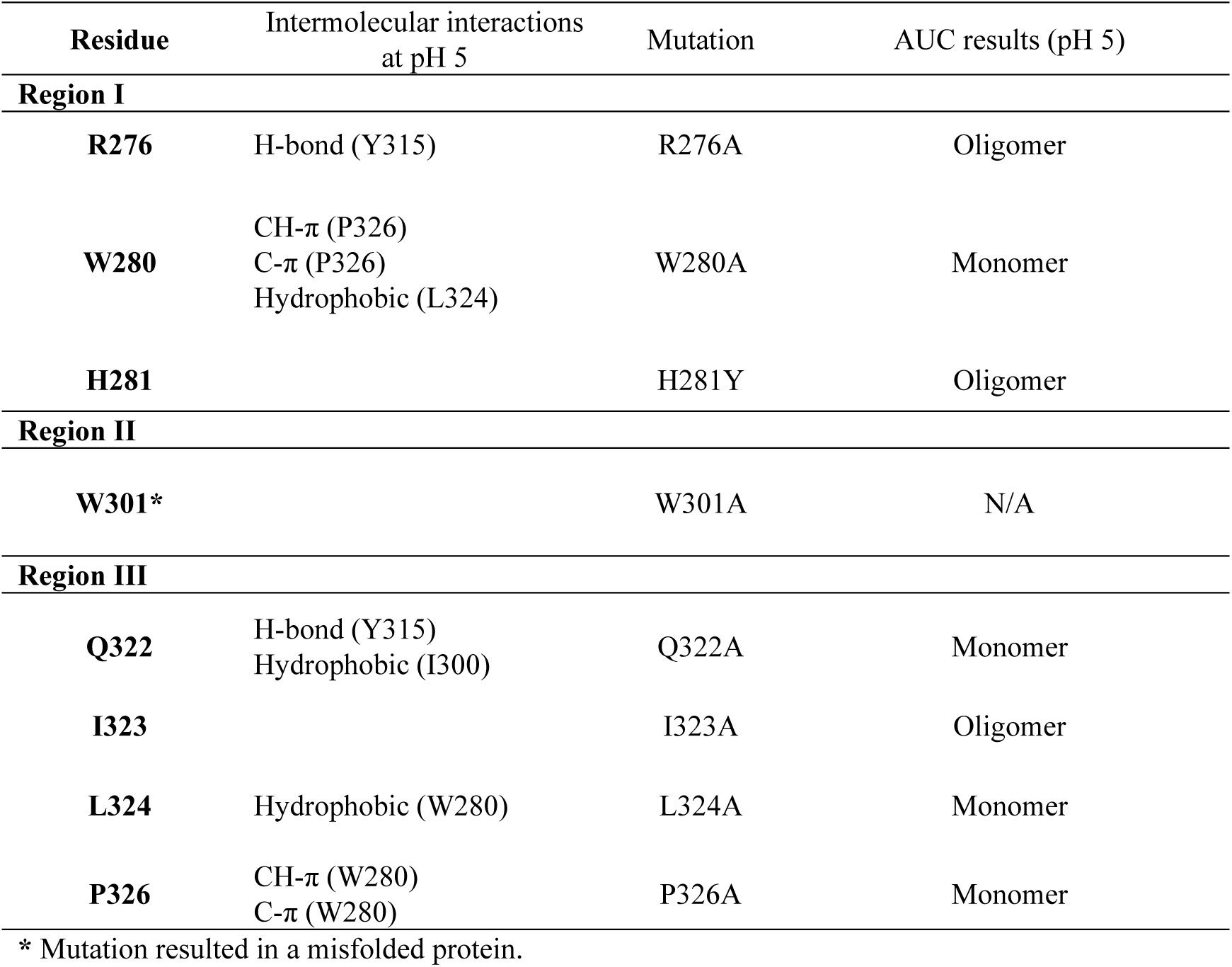
List of GNNV-P mutants.

| **Residue** | **Intermolecular interactions**<br>**at pH 5** | **Mutation** | **AUC results (pH 5)** |
| --- | --- | --- | --- |
| **Region I** |  |  |  |
| **R276** | H-bond (Y315) | R276A | Oligomer |
| **W280** | CH-π (P326)<br>C-π (P326)<br>Hydrophobic (L324) | W280A | Monomer |
| **H281** |  | H281Y | Oligomer |
| **Region II** |  |  |  |
| **W301\*** |  | W301A | N/A |
| **Region III** |  |  |  |
| **Q322** | H-bond (Y315)<br>Hydrophobic (I300) | Q322A | Monomer |
| **I323** |  | I323A | Oligomer |
| **L324** | Hydrophobic (W280) | L324A | Monomer |
| **P326** | CH-π (W280)<br>C-π (W280) | P326A | Monomer |

\* Mutation resulted in a misfolded protein.

### Amide hydrogen-deuterium exchange rate (HXD)

Hydrogen-deuterium exchange between amide NH signal and D_2_O solvent was monitored using ^1^H, ^15^N-HSQC spectra. First, an initial ^15^N-HSQC spectrum was collected before freezing GNNV-P in liquid nitrogen and lyophilizing it. The progress of amide hydrogen-deuterium exchange was monitored by collecting ^15^N-HSQC spectra for GNNV-P samples at pH 7.0 after re-dissolving them in 100% D_2_O for 4 hours. The hydrogen-deuterium exchange rate was calculated by fitting peak intensities into the exponential decay function I(t)=I_0·e^(-t/x) using NMRviewJ [53].

### Chemical shift perturbation and secondary chemical shifts calculation

2D ^1^H-^15^N HSQC spectra of GNNV-P at pH 7.0 and 5.0 were used for chemical shift perturbation analysis. The chemical shift between pH 7.0 and pH 5.0 was calculated using the equation 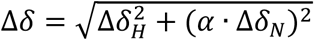, where Δδ_H_ and Δδ_N_ are the ^1^H and ^15^N chemical shift changes, respectively. A scaling factor of α = 0.1 was used to account for the larger ^15^N chemical shift [29]. Secondary structure propensities were predicted from ^13^C_α_ and ^13^C_β_ chemical shifts, based on their deviations from random coil values [59].

### Molecular dynamics (MD) simulation

Simulations for GNNV-P trimer formation were performed using the GROMACS package [60]. Initially, protein protonation state was adjusted to pH 5.0 using PDB2PQR software [61]. To prepare the initial point for the simulation, three protein molecules were initially packed in a ∼300 Å cubic box using the software PACKMOL [62]. This step was done to ensure that no repulsive interactions would disrupt or cause an error during the simulations. Using a V-rescale thermostat, the overall temperature of the water and protein were kept constant by coupling each group of molecules independently at 300 K. A Parrinello-Rahman barostat was used to separately couple the pressure to 1 atm in every dimension [63]. The time constants for the temperature and pressure coupling were set to 0.1 and 2 ps, respectively. A time step of 2 fs was applied using the leapfrog algorithm to integrate the equations of motion for the system. Periodic boundary conditions were set for the whole system. For the Lennard–Jones and the Ewald sum Coulombic interactions, we set a 1 nm cut-off. The Fourier space part of the Ewald splitting was calculated using the particle-mesh-Ewald method, by applying cubic spline interpolation and 0.16 nm grid length on the side [64]. The TIP3P water model was used and the protein parameters were obtained from the AMBERff99SB-ILDN force field [65,66]. MD simulations were done for a total scan length of 100 ns.

### Molecular docking analysis

Molecular docking analysis on GNNV-P with two sialoside isomers, Neu5Ac-(α2,3)-Lac and Neu5Ac-(α2,6)-Lac, was carried out using the HADDOCK 2.4 webserver and protein-glycan default settings [67]. The three-dimensional structures of Neu5Ac-(α2,3)-Lac and Neu5Ac-(α2,6)-Lac were extracted from PDB entries 6TLZ and 6TM0, respectively. The binding energy between GNNV-P and the docked sialosides was estimated using the PRODIGY webserver [68].

## Author Contributions

W.-H.C. conceived this project. C.-H.W. cultured and isolated native GNNV virus; expressed and purified GNNV virus-like particles (VLPs), and performed negative-stain electron microscopy as quality control. C.-H.W., T.K. and K.N. performed cryo-EM imaging on GNNV virus using cryo-ARM (JEOL Ltd., Akishima, Tokyo, Japan). C.-H.W. performed cryo-EM imaging on GNNV VLPs using JEM-2100F (JEOL Ltd., Akishima, Tokyo, Japan) and Technai F20 (FEI, Hillsboro, OR, USA). C.-H.W. performed all cryo-EM image analysis. P.S. and C.-H.W. performed cloning of the GNNV P-domain. P.S. performed expression and purification of the P-domain, followed by analytical ultracentrifugation (AUC) analysis. P.S., Y.-C.L., and D.-L.M.T. performed NMR experiments and respective analysis. P.S. and C.-H.W. performed P-domain mutagenesis, and P.S. conducted AUC tests for the mutants. K.J.D.C. performed molecular dynamic simulations. P.S. performed molecular docking analysis. P.S., C.-H.W., D.-L.M.T., and W-H.C. prepared the manuscript.

## Acknowledgements

This project was supported by MOST grants [103-2321-B-001-048], [104-2321-B-001-019], [105-2321-B-001-009], and AS SUMMIT Projects [AS-SUMMIT-106] and [AS-SUMMIT-107] to W.-H.C, and MOST grant [107-2113-M-001-017] to D.-L. M. T. The authors are indebted to Yeukuang Hwu and Yi-Yun Chen at the Institute of Physics, Academia Sinica, for supporting the use of JEM-2100F (JEOL Ltd., Akishima, Tokyo, Japan), as well as to Dong-Hua Chen, David Bushnell and Roger Kornberg at Stanford University for supporting the use of Technai F20 (FEI, Hillsboro, OR, USA). The operation of cryo-ARM at Osaka University was supported by a Japan Joint Research Grant to K.N. Note that all of the cryo-EM data for this work were obtained prior to installation of the Academia Sinica Cryo-EM Core Facility (ASCEM) supported by [AS-CFII-108-110]. The authors thank the Academia Sinica High-Field NMR Center (HFNMRC; funded by Academia Sinica Core Facility and Innovative Instrument Project AS-CFII-111-214) for technical support. The authors thank the Medicinal Chemistry and Analytical Core Facilities, funded by Academia Sinica Core Facility and Innovative Instrument Project AS-NBRPCF-111-201, for technical support with NMR data. We thank Mr. Kun Hung Chen in the Biophysics Core Facility, funded by Academia Sinica Core Facility and Innovative Instrument Project AS-CFII-111-201, for performing the sedimentation velocity analytical ultracentrifugation experiments. The authors thank the DNA Sequencing Core Facility of the Institute of Biomedical Sciences, Academia Sinica (funded by Academia Sinica Core Facility and Innovative Instrument Project AS-CFII-111-211), for DNA sequencing analysis. This study made use of NMRbox: National Center for Biomolecular NMR Data Processing and Analysis, a Biomedical Technology Research Resource (BTRR), which is supported by NIH grant P41GM111135 (NIGMS).

## Data Availability

Cryo-EM maps of GNNV VLP at pH 8.0, 6.5 and 5.0 are available from the EMDB data bank with accession numbers EMDB-39212, 39213, and 39214, and the corresponding atomic models are with PDB entries of 8YF6, 8YF7, and 8YF8 respectively; Cryo-EM maps of GNNV virion at pH 6.5 and 5.0 have EMDB accession numbers 39215 and 39217, and the atomic model of GNNV virion at pH 6.5 is with PDB entry of 8YF9. The chemical shift assignments for 2D HSQC spectra at pH 5.0 have been deposited to the Biological Magnetic Resonance Bank (BMRB) with Accession No. 52218. The NMR structure of the GNNV P-domain has been deposited to the Protein Data Bank (PDB entry 8XID).

