## Supplementary Information for "Cryo-EM inspired NMR analysis reveals a pH-induced conformational switching mechanism for imparting dynamics to Betanodavirus protrusions"

### for

**This file contains 21 supplementary figures and 3 supplementary movies.**

**Fig. S1. Cryo-EM images of GNNV in different pH environments.**

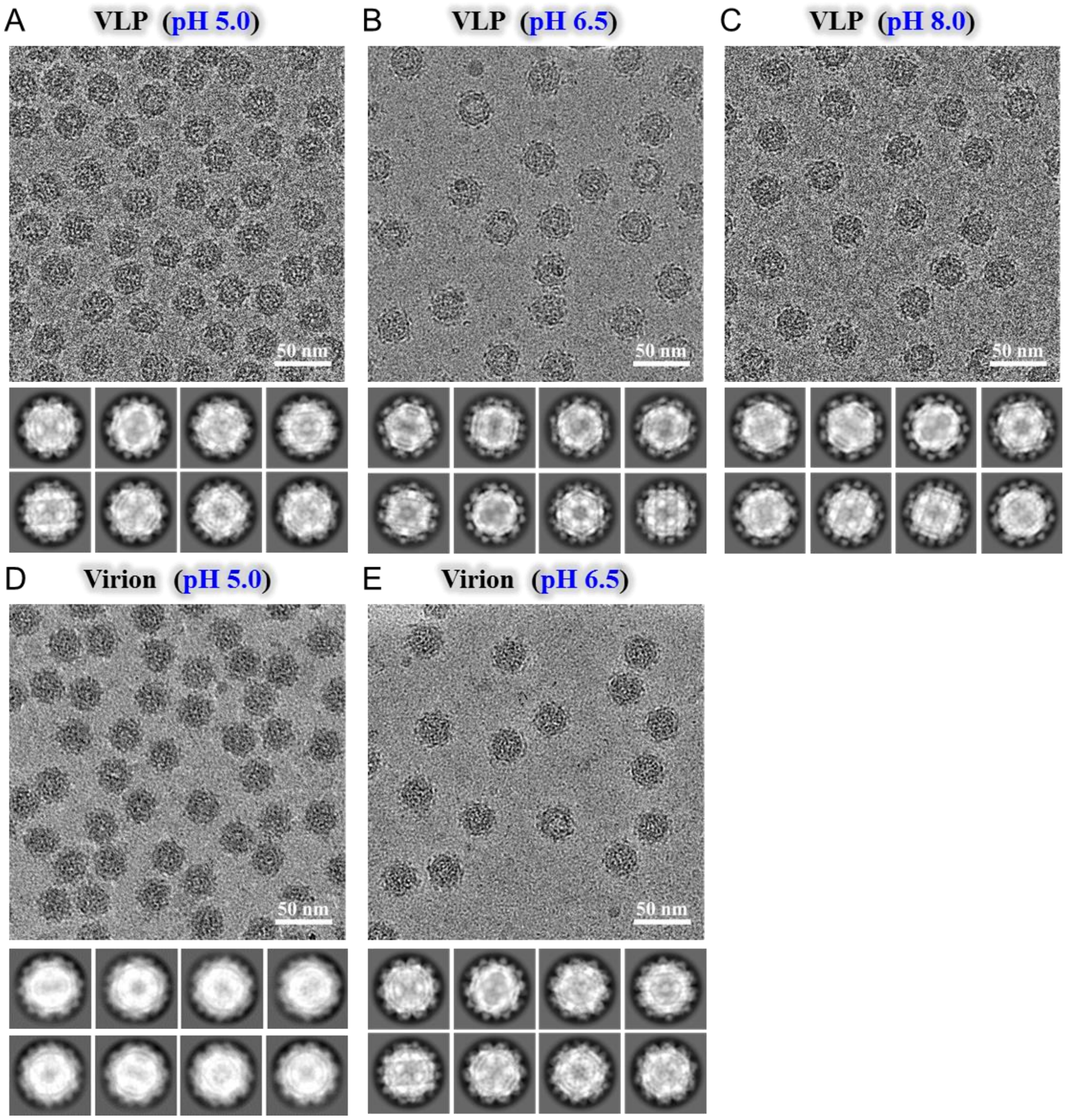


**Supplementary Figure 1. Cryo-EM micrographs class averages of GNNV in different pH environments.** **(A)** GNNV VLP at pH 5.0. bar: 50 nm. **(B)** GNNV VLP at pH 6.5. bar: 50 nm. **(C)** GNNV VLP at pH 8.0. bar: 50 nm. **(D)** GNNV virion at pH 5.0. bar: 50 nm. **(E)** GNNV virion at pH 6.5. bar: 50 nm.

**Fig. S2. Structure of native GNNV virions compared to virus-like particles (VLP) at the same pH.**

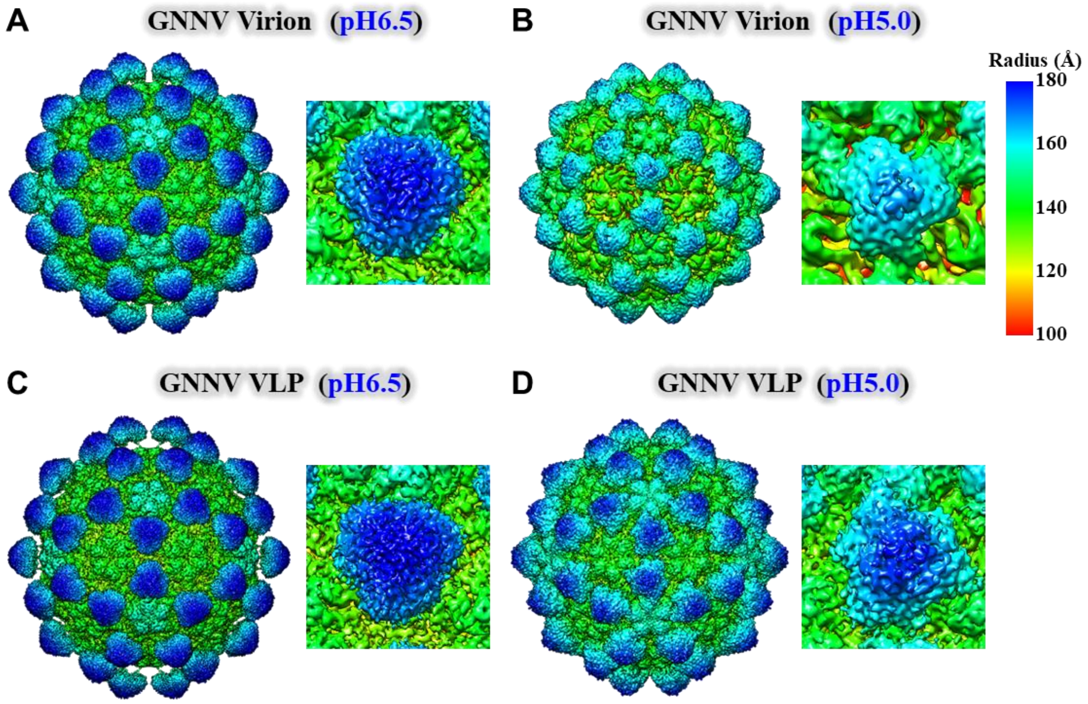


**Supplementary Figure 2. Structure of native GNNV virions compared to virus-like particles (VLP) at the same pH. (A)** GNNV virion at pH 6.5 (resolution 3.12 Å, contours at 2σ). **(B)** GNNV virion at pH 5.0 (resolution 4.36 Å, contours at 3σ). **(C)** GNNV VLP at pH 6.5 (resolution 2.72 Å, contours 2σ). **(D)** GNNV VLP at pH 5.0 (resolution 3.65 Å, contours at 2σ). The structure of GNNV is colored from red to blue according to the radius, as shown by the color bar. In the panels at right of A-D, a magnified view of a protrusion is shown. Note that the VLP data at pH 6.5 was collected using an F20 electron microscope (FEI, Hillsboro, OR, USA) with a K2 camera (Gatan Inc., Pleasanton, CA, USA), whereas the VLP data at pH 5.0 was collected using a JEM-2100F electron microscope (JEOL Ltd., Akishima, Tokyo, Japan) with a DE-20 camera (Direct Electron LP, San Diego, CA, USA).

**Fig. S3. Cryo-EM structural determination of GNNV VLPs at pH 8.0, pH 6.5 and pH 5.0.**

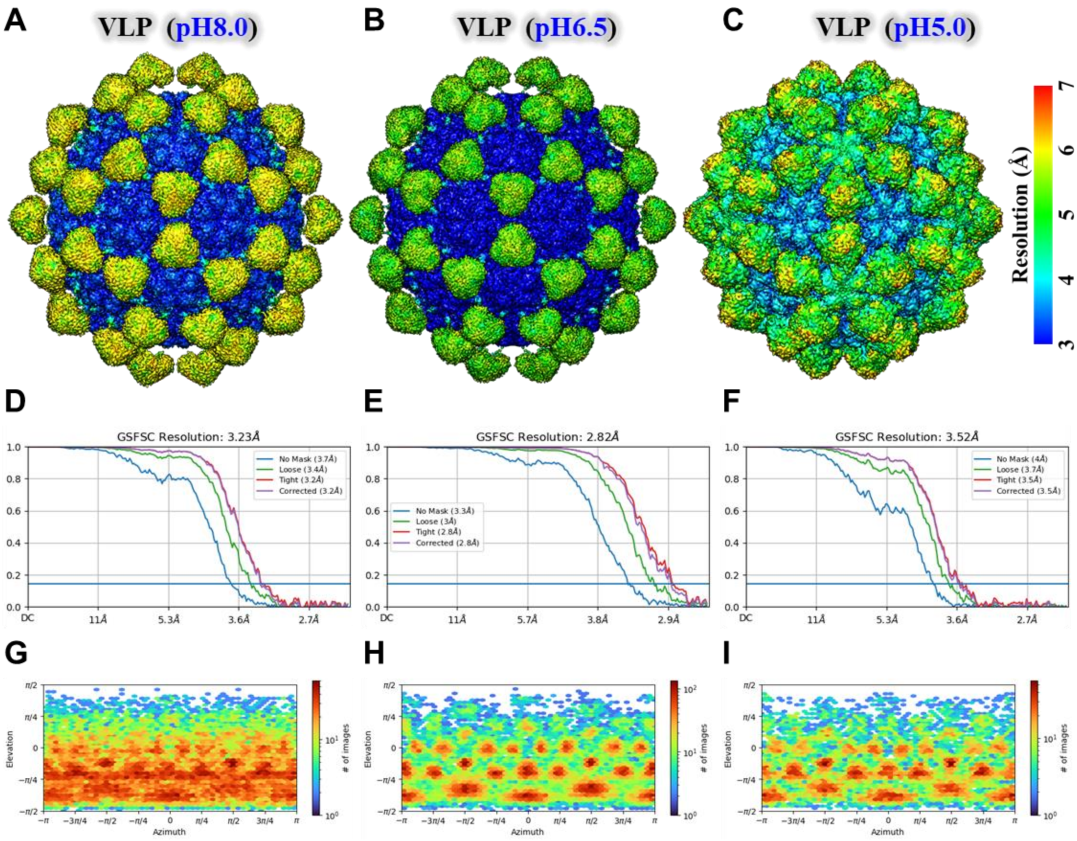


**Supplementary Figure 3. Cryo-EM structural determination of GNNV VLPs at pH 8.0, pH 6.5 and pH 5.0.** Local resolution analysis of the cryo-EM maps of GNNV VLPs at **(A)** pH 8.0, **(B)** pH 6.5, and **(C)** pH 5.0. The cryo-EM maps of GNNV VLPs are colored according to the resolution, as shown by the color bar. Gold-standard FSC curves (FSC = 0.143) of the cryo-EM map of GNNV VLPs at **(D)** pH 8.0, **(E)** pH 6.5, and **(F)** pH 5.0. The angular distributions of all particle projections in the final 3D reconstructions of GNNV VLPs at **(G)** pH 8.0, **(H)** pH 6.5, and **(I)** pH 5.0. The heat maps depict the number of particles observed for each viewing angle. Regions colored in red indicate a higher particle count, indicating a more frequent occurrence of those specific viewing angles.

**Fig. S4. Cryo-EM structural determination of GNNV virions at pH 6.5 and pH 5.0.**

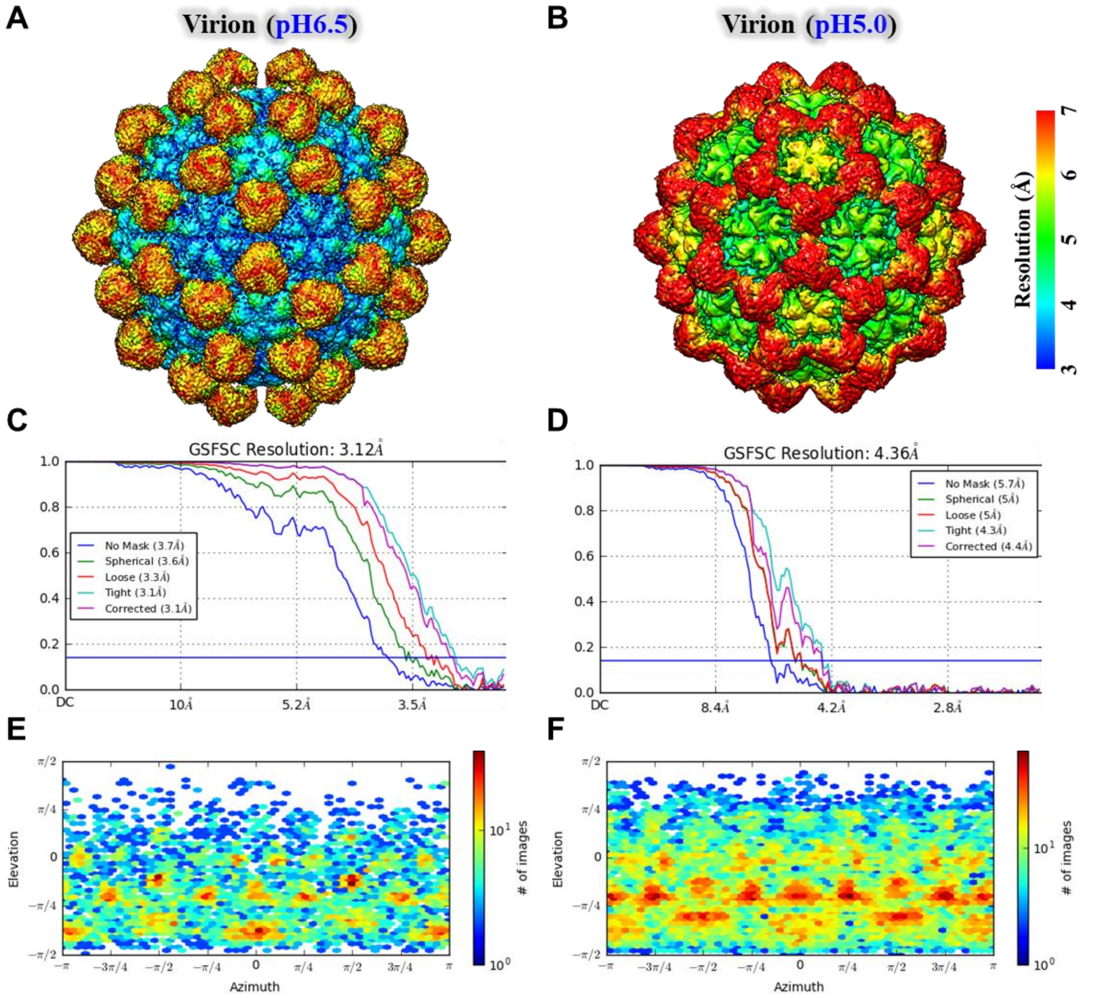


**Supplementary Figure 4. Cryo-EM structural determination of GNNV virions at pH 6.5 and pH 5.0.** Local resolution analysis of the cryo-EM maps of GNNV virions at **(A)** pH 6.5 and **(B)** pH 5.0. The cryo-EM maps of GNNV virions are colored according to the resolution, as shown by the color bar. Gold-standard FSC curves (FSC = 0.143) of the cryo-EM maps of GNNV virions at **(C)** pH 6.5 and **(D)** pH 5.0. The angular distributions of all particle projections in the final 3D reconstructions of GNNV virions at **(E)** pH 6.5 and **(F)** pH 5.0. The heat maps depict the number of particles observed for each viewing angle. Regions colored in red indicate a higher particle count, indicating a more frequent occurrence of those specific viewing angles.

**Fig. S5. Particle diameters of GNNV VLPs at different pH and the radial density distribution of GNNV VLPs.**

A

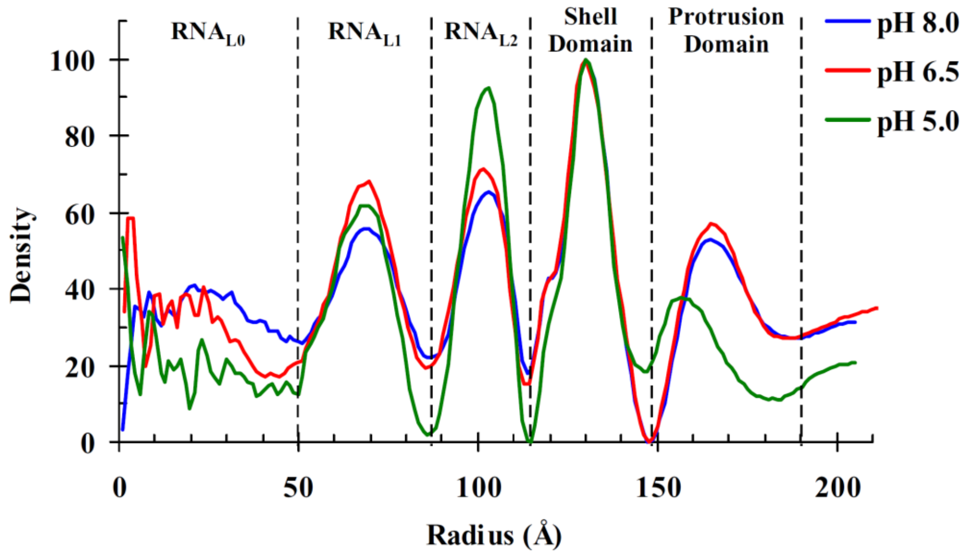


**Supplementary Figure 5. Particle diameters of GNNV at different pH and radial density distribution of GNNV. (A)** The surface view of GNNV particles at pH8.0 (*left panel*), pH6.5 (*middle panel*) and pH5.0 (*right panel*). The maps are colored based on the radius. **(B)** The radial density distribution of GNNV VLPs in various pH environments. Three sections correspond to the protrusions (~149 to 190 Å), the capsid shell (~115 to 149 Å).

**Fig. S6. Cross-sections of GNNV VLPs in three different pH environments.**

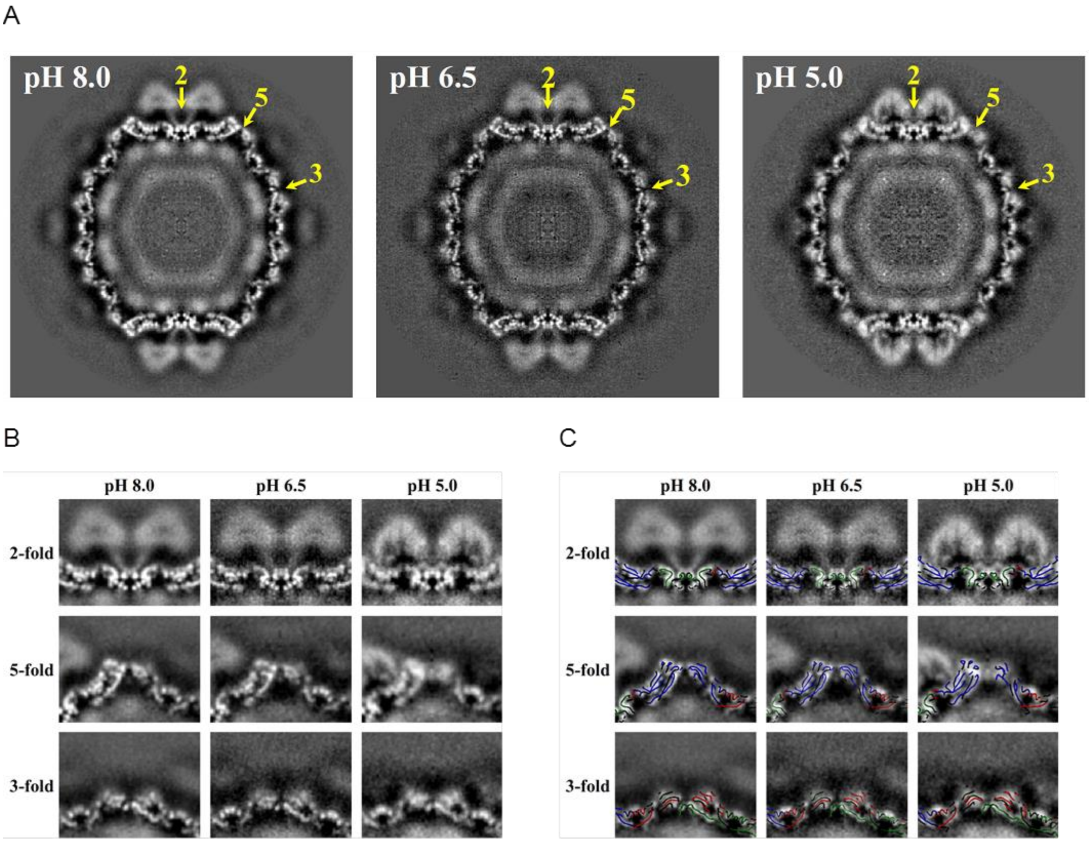


**Supplementary Figure 6. Cross-sections of GNNV VLPs in three different pH environments. (A)** Cross-sections of a GNNV particle at pH 8.0 (*left panel*), pH 6.5 (*middle panel*), and pH 5.0 (*right panel*). Icosahedral 2-, 5- and 3-fold axes are indicated. **(B)** Enlarged views of the cross-sections around the icosahedral 2-, 5- and 3-fold axes in (A). **(C)** Fitting of atomic models of the S-domain subunits A, B, and C (colored blue, red and green, respectively) into the cross sections in (B).

**Fig. S7. NMR spectra of deuterium-labeled GNNV-P at pH 5.0.**

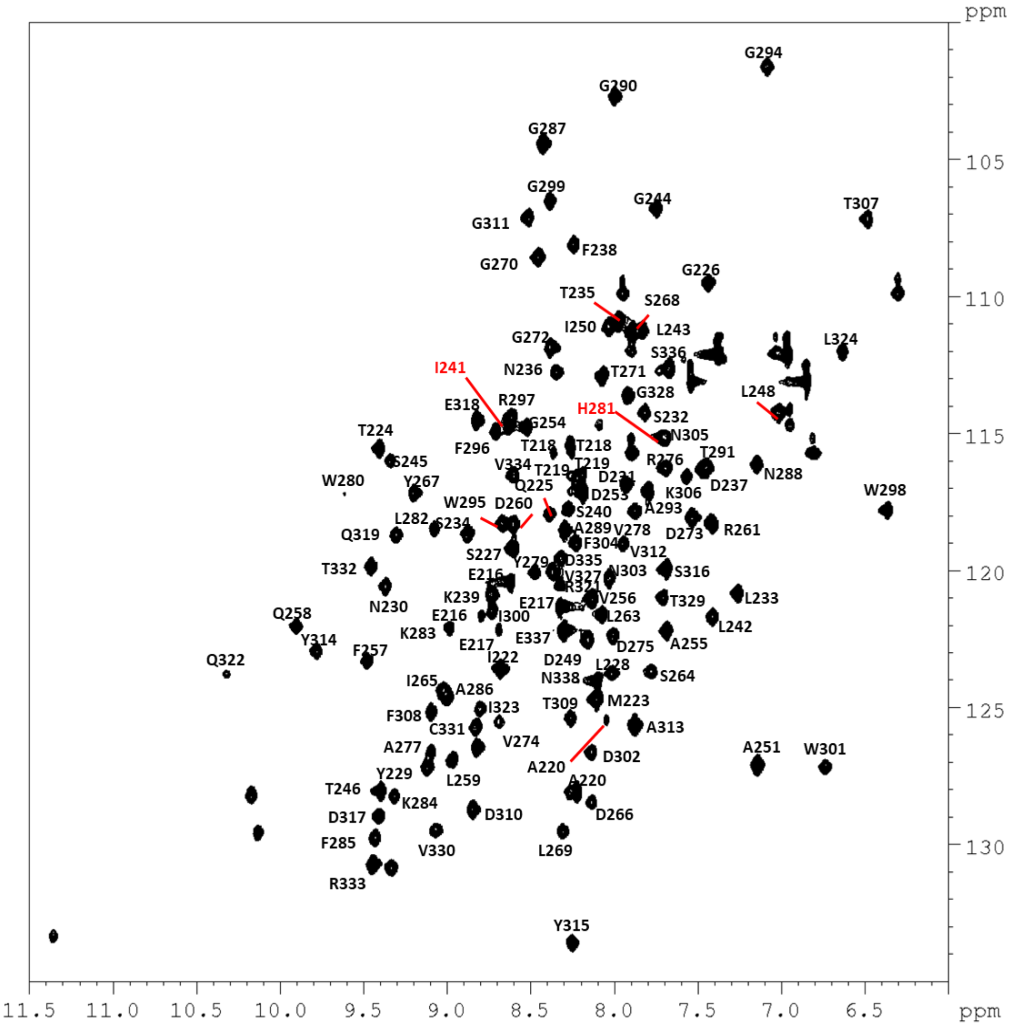


**Supplementary Figure 7. NMR spectra of deuterium-labeled GNNV-P at pH 5.0.** HSQC spectra of $^{2}$H-, $^{13}$C-, and $^{15}$N-labeled GNNV-P recorded at pH 5.0. The resonance assignments for pH 5.0 have been deposited into the Biological Magnetic Resonance Databank with Accession No. 52218.

**Fig. S8. NMR spectra reveal the effects of pH on GNNV-P**

A

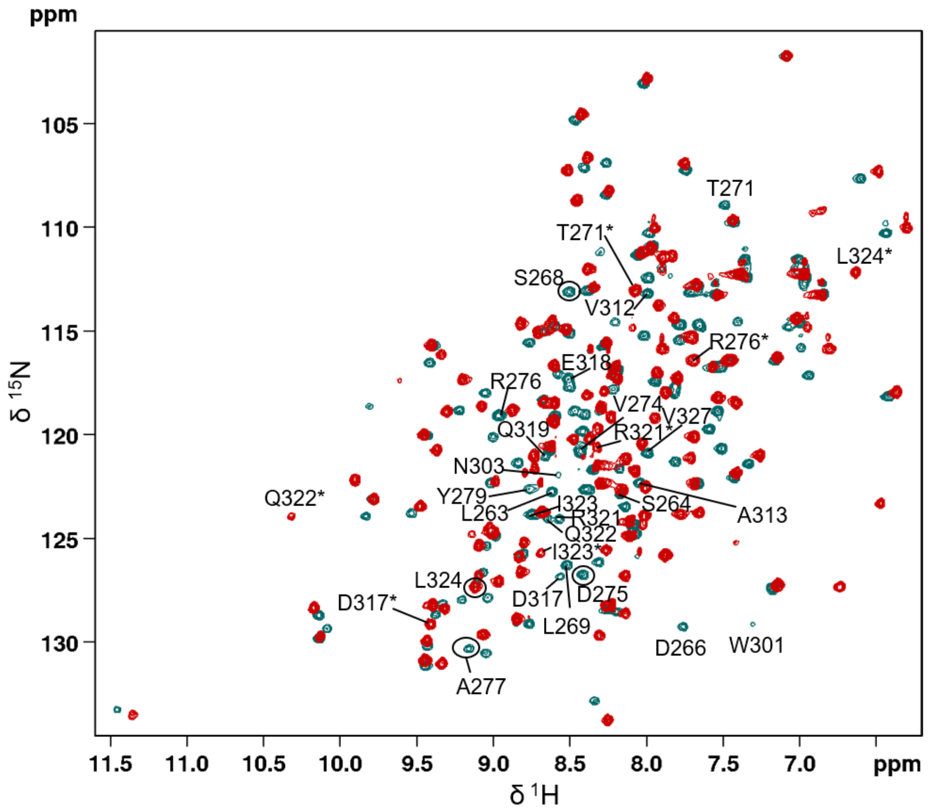


B

TPEETTAPIMTQGSLYNDSLSTNDFKSILLGSTPLDIAPD 40
GAVFQLDRPLSIDYSLGTGDVDRAVYWHLKKFAGNAGTPA 80
GWFRWGIWDNFNKTFTDGVAYYSDEQPRQILLPVGTVCTR 120
VDSEN 125

**Supplementary Figure 8. NMR spectra reveal the effects of pH on GNNV-P. (A)** Superimposition of $^{1}H$-$^{15}N$ HSQC spectra of $^{15}N$-labeled GNNV-P recorded at pH 7.0 (blue) and $^{2}H$-, $^{13}C$-, and $^{15}N$-labeled GNNV-P recorded at pH 5.0 (red). Resonances of the pH-sensitive peaks at pH 7.0 are labeled. New positions for residues with the greatest chemical shift perturbations (CSPs) at pH 5.0 are marked by *. **(B)** GNNV-P amino acid sequence with pH-sensitive residues highlighted in blue. Red residues represent those displaying the greatest CSPs between neutral and acidic pH.

**Fig. S9. Sedimentation velocity analytical ultracentrifugation analysis of GNNV-P oligomerization.**

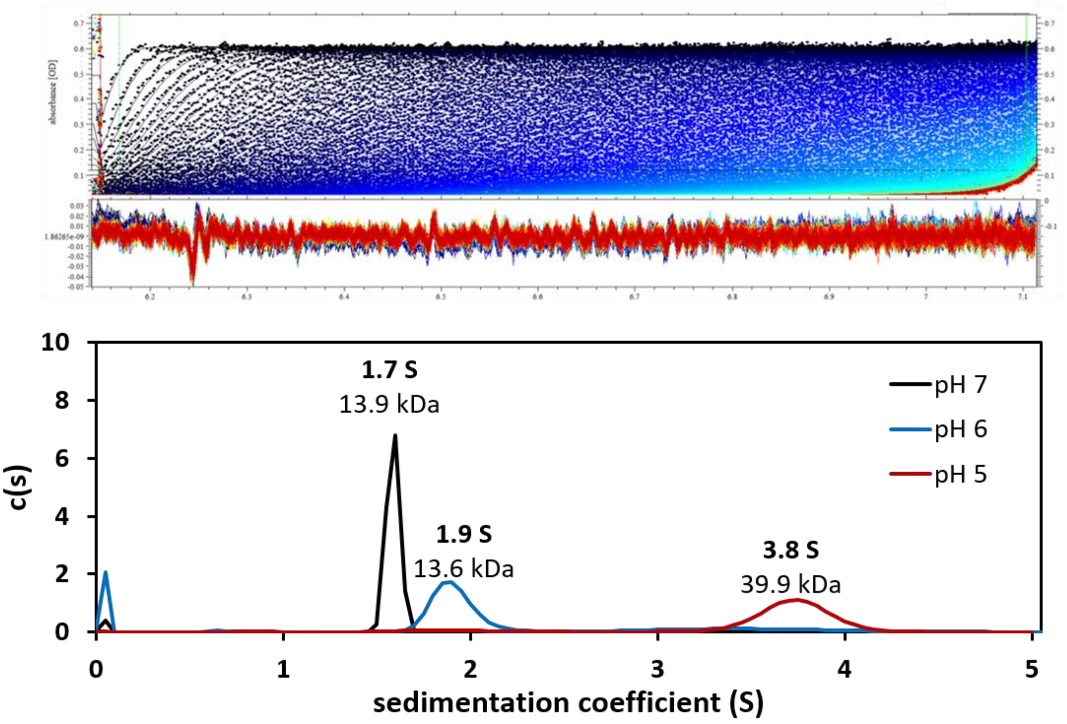


**Supplementary Figure 9. Sedimentation velocity analytical ultracentrifugation analysis of GNNV-P oligomerization.** Plot of the sedimentation velocity analytical ultracentrifugation (SV AUC) data fitted to a continuous sedimentation coefficient distribution c(s) model. The GNNV-P sedimentation coefficients were determined at pH 7.0 (black), 6.0 (blue), and 5.0 (red). Sedimentation coefficients and estimated molecular weights are noted above each peak. Representative raw sedimentation profile of absorbance at 280 nm and representative residuals from fitting the data to a continuous c(s) distribution model are shown in the panels above the plot.

**Fig. S10. Hydrogen-deuterium exchange rate (HXD) of GNNV-P at pH 7.0.**

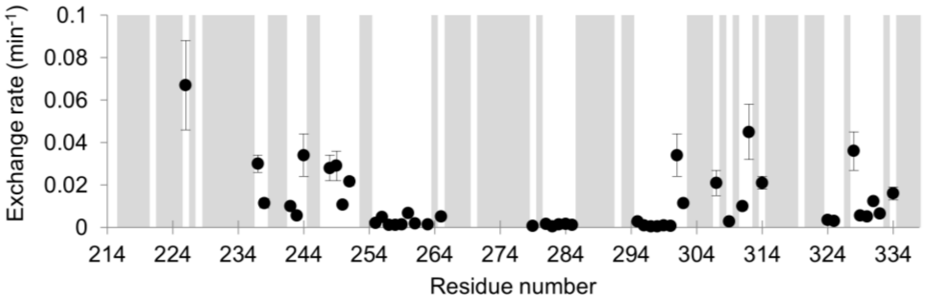


**Supplementary Figure 10. Hydrogen-deuterium exchange rate (HXD) of GNNV-P at pH 7.0.** Plot of the hydrogen-deuterium exchange rate determined at neutral pH as a function of protein sequence. Residues for which HXD could not be measured (i.e., exchanging faster than our detection limits) are indicated by a grey background.

**Fig. S11. Mapping of pH-sensitive residues on the GNNV-P trimer crystal structure.**

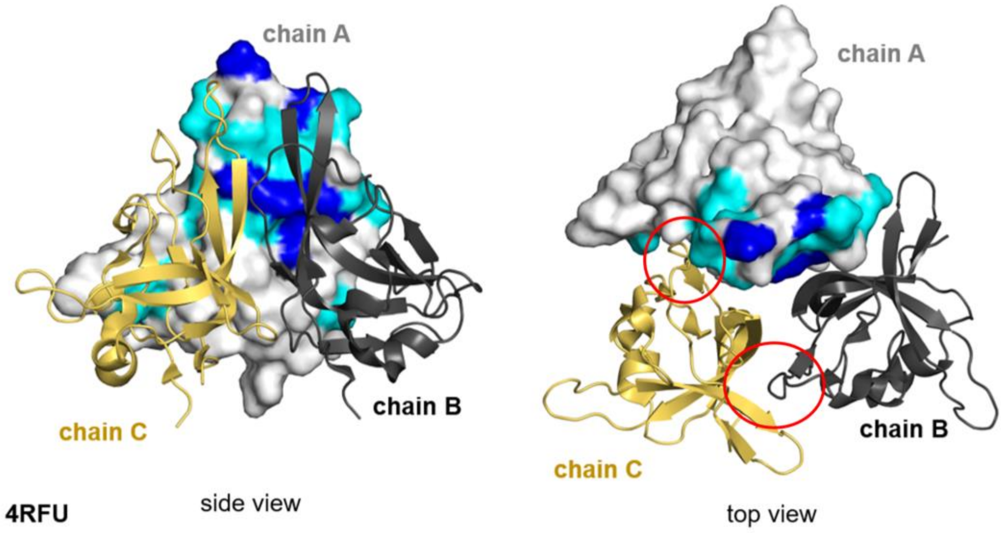


**Supplementary Figure 11. Mapping of pH-sensitive residues on the GNNV-P trimer crystal structure.** pH-sensitive residues identified by NMR pH titration mapped onto the crystal structure of trimeric GNNV-P (PDB ID 4RFU). The pH-sensitive residues have been mapped onto the surface representation of chain A consistent with the color scheme in Fig. 2C. Chain B and chain C are displayed as yellow and black cartoons, respectively. The F'-G' loop and β-strand region (aa 323-326) at the trimeric interface are highlighted by red circles.

**Fig. S12. RMSD analysis of GNNV-P trimer stability after MD simulations.**

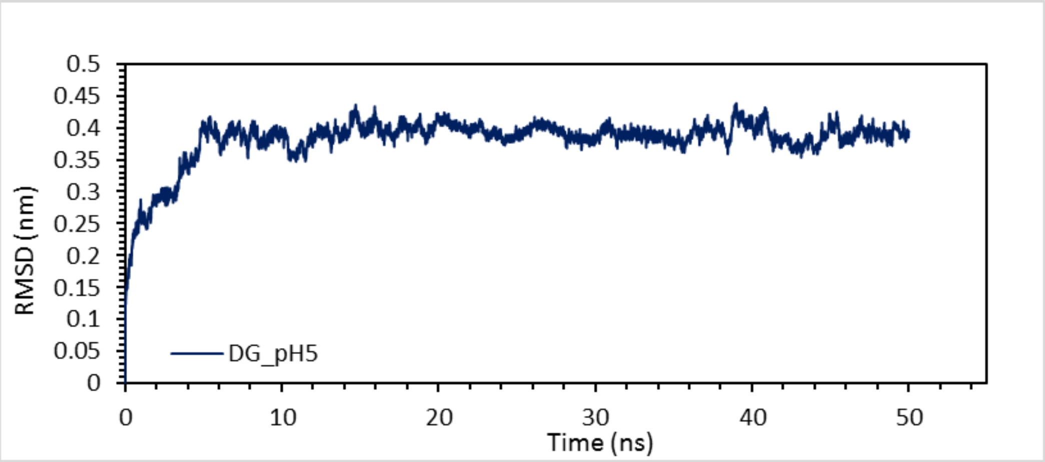


**Supplementary Figure 12. RMSD analysis of GNNV-P trimer stability after MD simulations.** RMSD of GNNV-P trimer during the time-course of MD simulations.

**Fig. S13. Comparison of the GNNV-P trimers obtained by MD simulations and X–ray crystallography.**

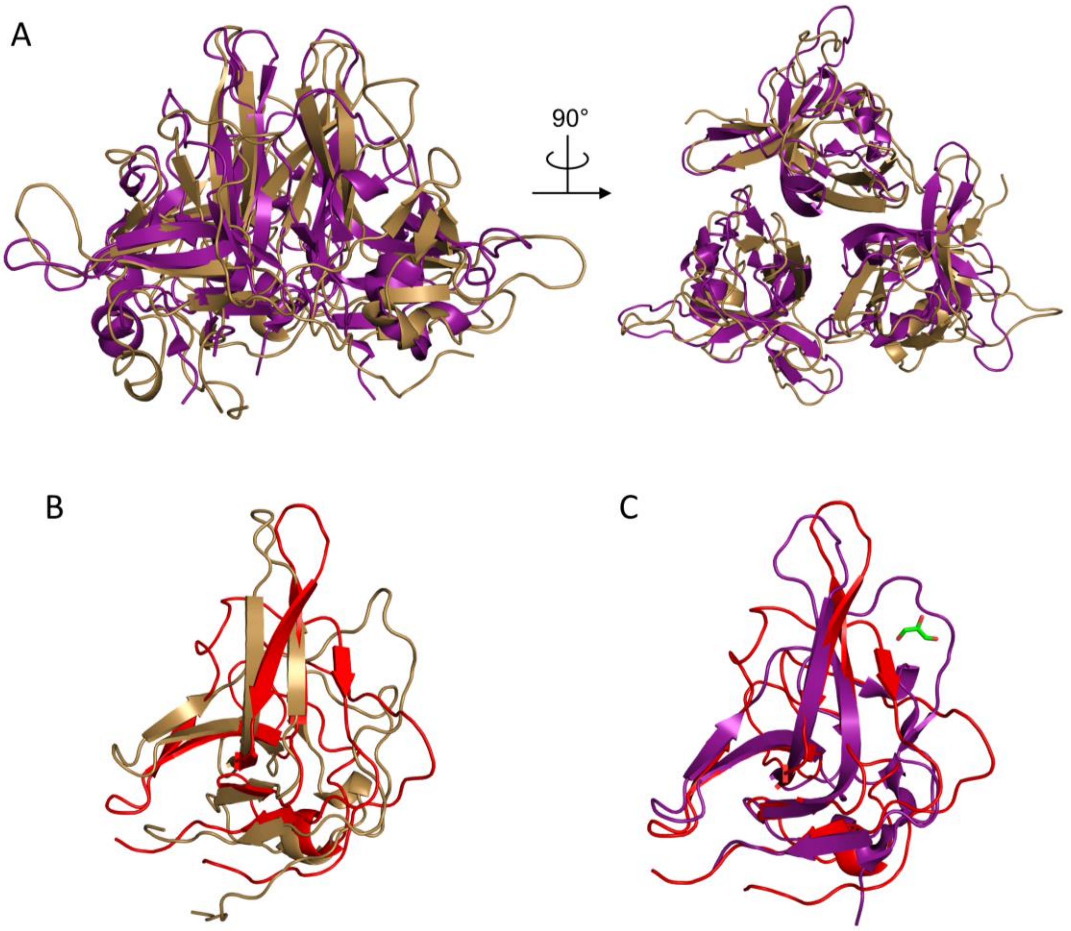


**Supplementary Figure 13. Comparison of the GNNV-P trimers obtained by MD simulation and X–ray crystallography.** (A) Side and top views of overlaid cartoon representations of GNNV-P trimers determined by MD simulations at pH 5.0 (purple) and X-ray crystallography at pH 6.5 (brown). (B) Superimposition of the GNNV-P structures determined in solution at neutral pH (light brown) and the model of GNNV-P at pH 5.0 obtained by MD simulations (red). (C) Superimposition of the MD-simulated GNNV-P model at pH 5.0 (red) with chain A of the GNNV-P structure determined by X-ray crystallography at pH 6.5 (purple). A conserved glycerol molecule resolved as stabilizing the F'-G' loop is shown in green as a stick model.

**Fig. S14. Sedimentation velocity AUC analysis of low-pH-induced oligomerization of GNNV-P single mutants.**

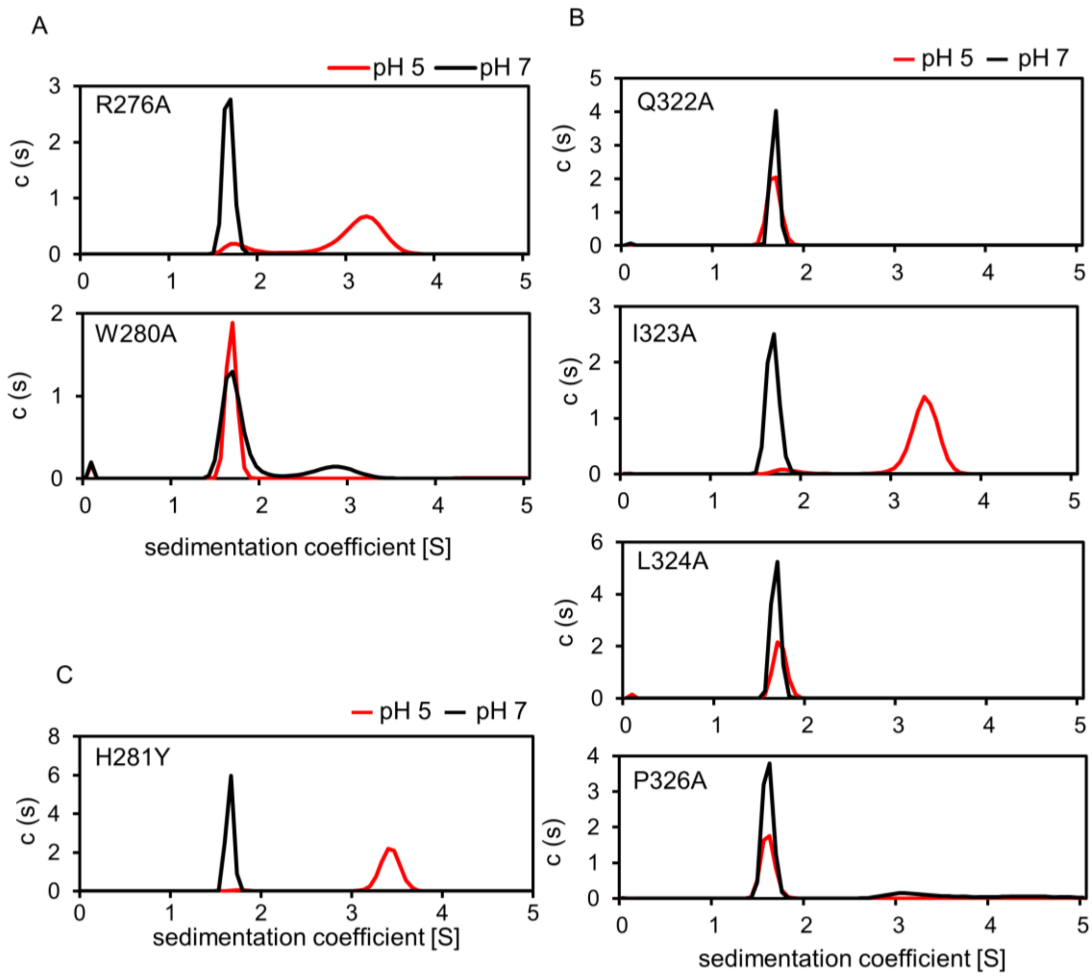


**Supplementary Figure 14. Sedimentation velocity AUC analysis of low-pH-induced oligomerization of GNNV-P single mutants.** Plots of the sedimentation velocity analytical ultracentrifugation (SV AUC) data fitted to a continuous sedimentation coefficient distribution c(s) model for GNNV-P mutations of residues in Region I (A), Region III (B), and for residue H281 (C).

**Fig. S15. Comparative protein sequence analysis of NNV P-domains.**

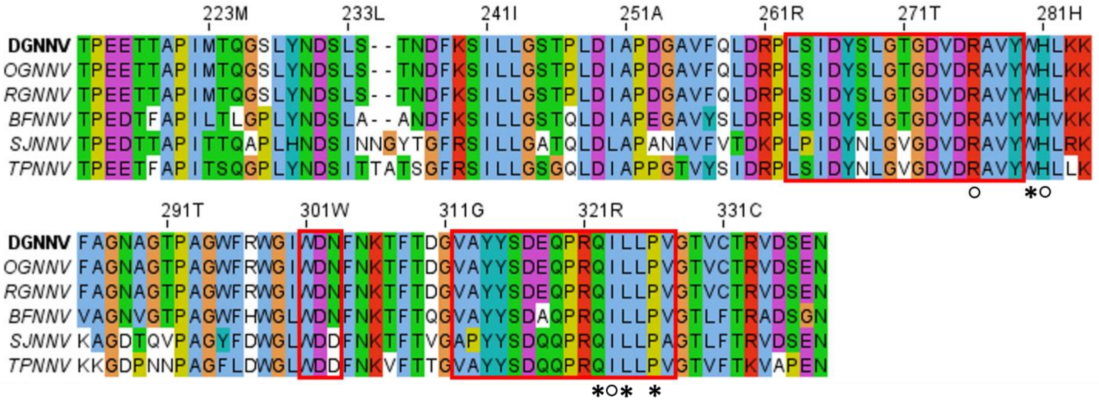


**Supplementary Figure 15. Comparative protein sequence analysis of NNV P-domains.** Protein sequence alignment of the P-domains of Dragon grouper nervous necrosis virus (DGNNV), Orange-spotted grouper nervous necrosis virus (OGNNV), plus four genotypic variants (RGNNV, BFNNV, SJNNV, and TPNNV), shows a high degree of conservation of amino acid residues within the F'-G' loop. pH-sensitive region I (L263-Y279), region II (W301-N303), and region III (V312-V327) are highlighted by red boxes. Residues identified by mutagenesis as being critical for GNNV-P low-pH-induced oligomerization are marked by *****. Residues for which mutation did not affect low-pH-induced oligomerization are marked by ○.

**Fig. S16. Changes in the electrostatic surface potential of monomeric GNNV-P at neutral and low pH.**

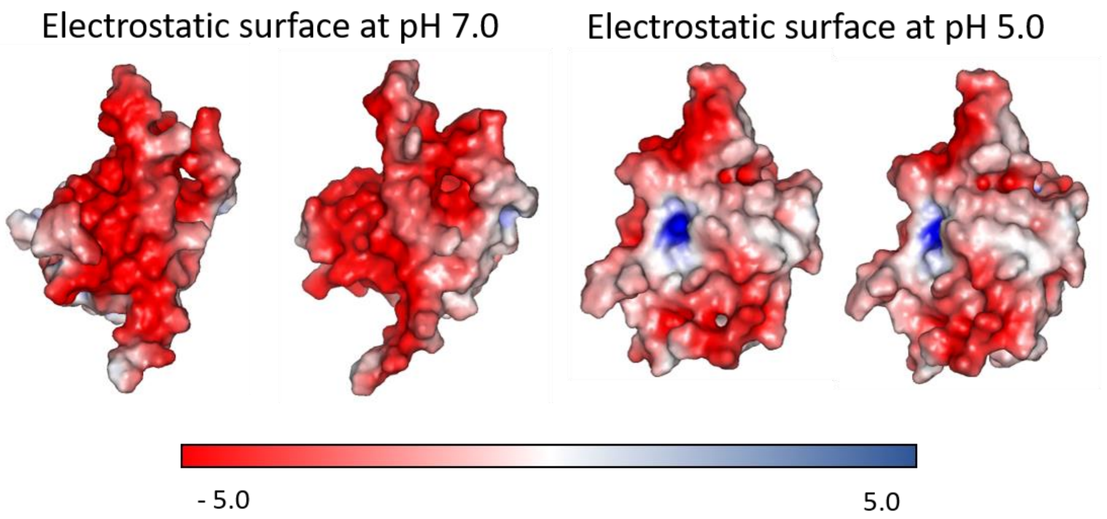


**Supplementary Figure 16. Changes in the electrostatic surface potential of monomeric GNNV-P at neutral and low pH.** Gradient visualization of the electrostatic surfaces from red (-5.0 kT/e) to blue (5.0 kT/e) of GNNV-P at pH 7.0 and 5.0, calculated using the APBS server based on residual pKa determined by PROPKA. The view shows the potential interface area between the A/C (left) and A/B (right) units in the GNNV-P trimer at pH 7.0 and pH 5.0.

**Fig. S17. The conformational change of the GNNV-P malleable linker between neutral and low pH.**

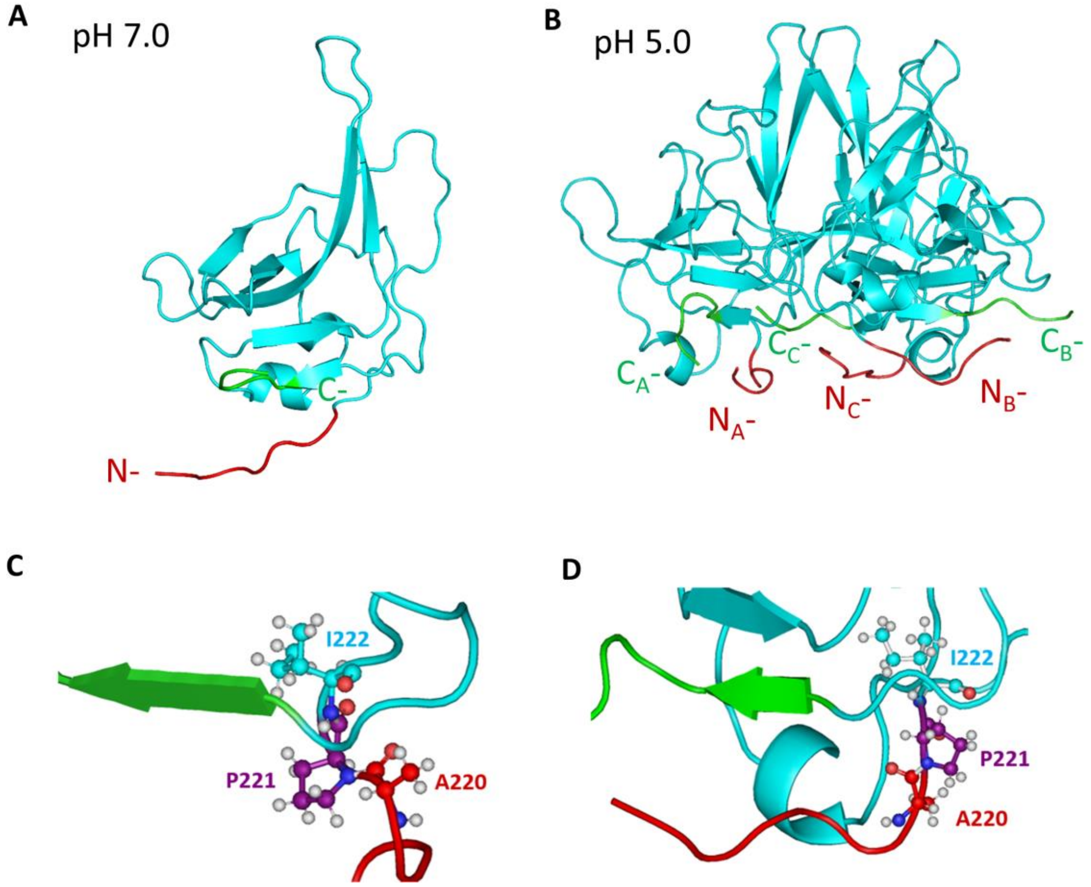


**Supplementary Figure 17. The conformational change of the GNNV-P malleable linker between neutral and low pH.** GNNV-P structure at neutral pH (A) and model of the GNNV-P trimer at acidic pH (B) with the N- and C-terminal residues colored red and green, respectively. Detailed view of the conformation of the malleable linker around residue P221 at neutral pH (C) and acidic pH (D). Residues A220-I222 are shown in ball and stick representation.

**Fig. S18. Multiple NMR assignments of residue A220 at low pH.**

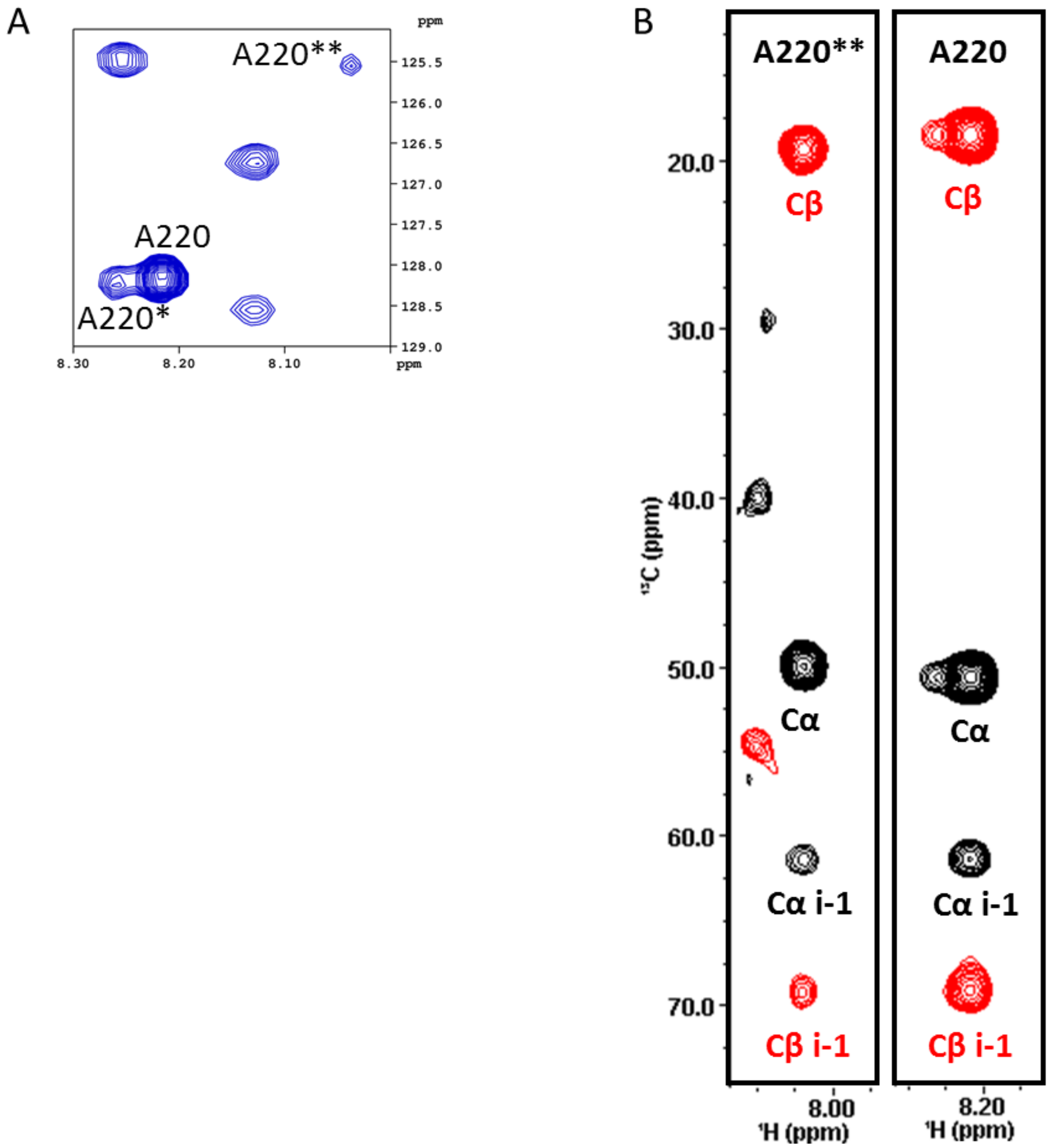


**Supplementary Figure 18. Multiple NMR assignments of residue A220 at low pH. (A)** Zoomed in view of the 2D 1H-15N HSQC spectrum of GNNV-P at pH 5.0, showing multiple NMR signals for residue A220. **(B)** HNCACB spectra showing the chemical shift of residue A220.

**Fig. S19. Model of Neu5Ac-Lac binding to GNNV-P at pH 5.0.**

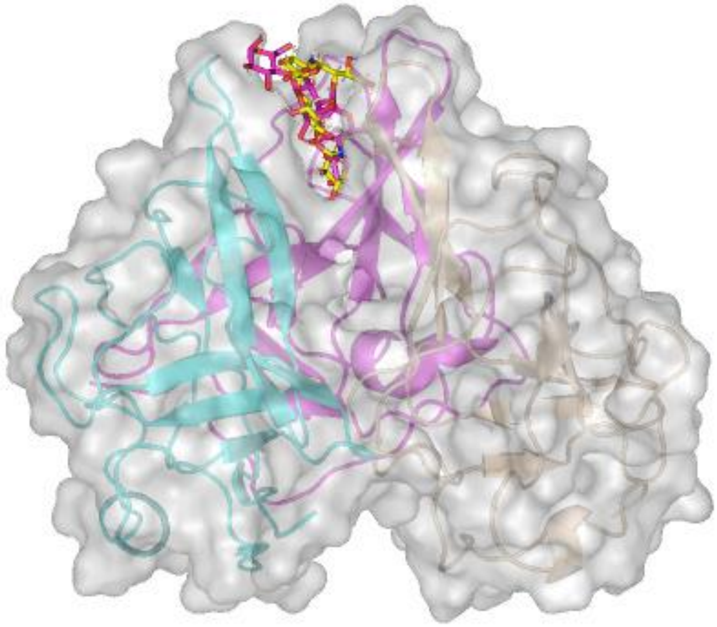

**Supplementary Figure 19. Model of Neu5Ac-Lac binding to GNNV-P at pH 5.0.** Models of Neu5Ac-(α2,3)-Lac (magenta) and Neu5Ac-(α2,6)-Lac (yellow) binding to the GNNV-P trimer (shown as cartoon and surface representations).

**Fig. S20. Flowchart of GNNV VLP image processing.**

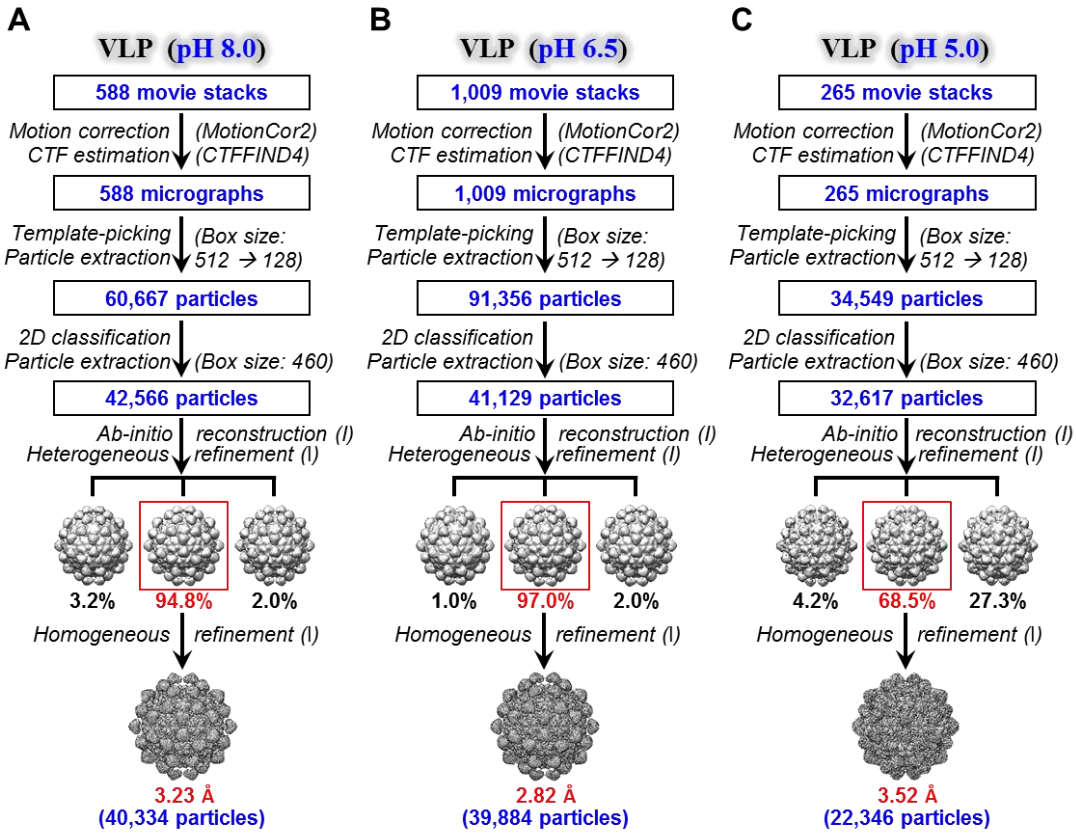


**Supplementary Figure 20. Flowchart of GNNV VLP image processing. (A)** GNNV VLP at pH 8.0. **(B)** GNNV VLP at pH 6.5. **(C)** GNNV VLP at pH 5.0.

**Fig. S21. Flowchart of GNNV virion image processing.**

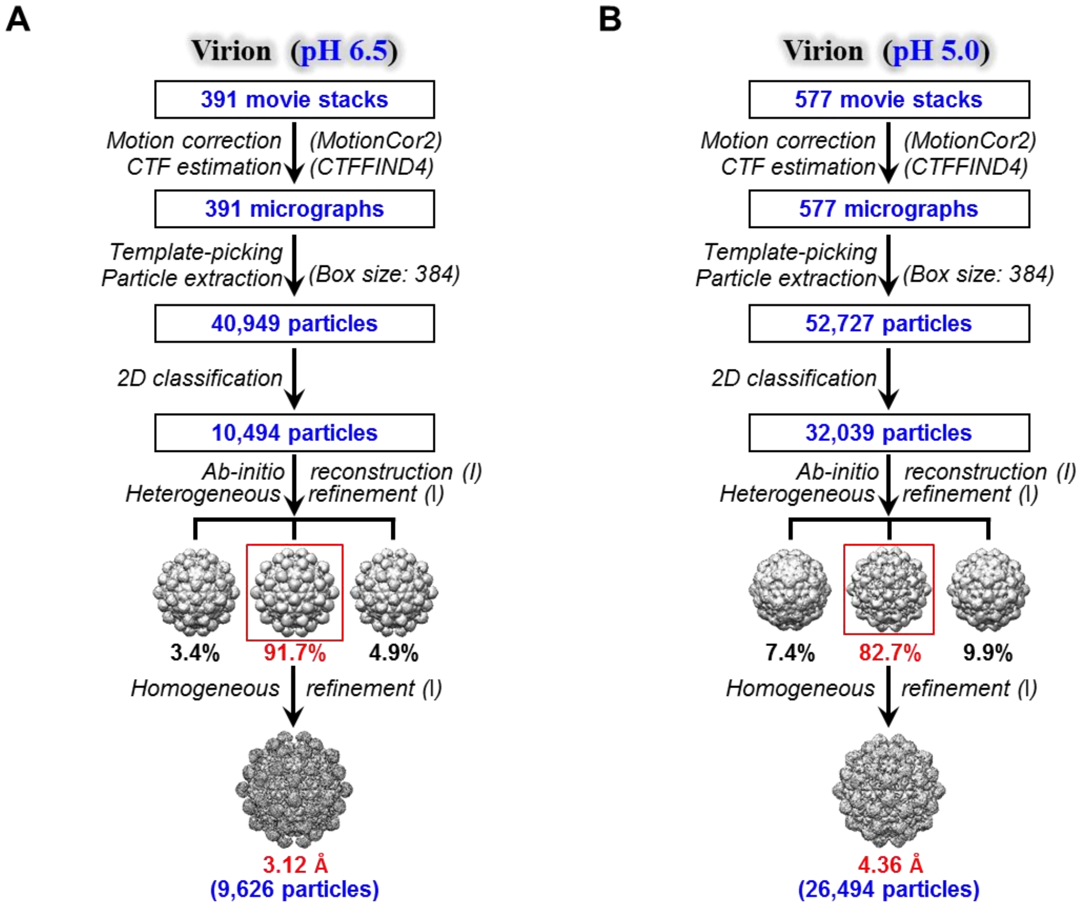


**Supplementary Figure 21. Flowchart of GNNV virion image processing. (A)** GNNV virion at pH 6.5. **(B)** GNNV virion at pH 5.0.

**Movie S1. Conformational change of GNNV VLP from pH 8.0 to 5.0.**

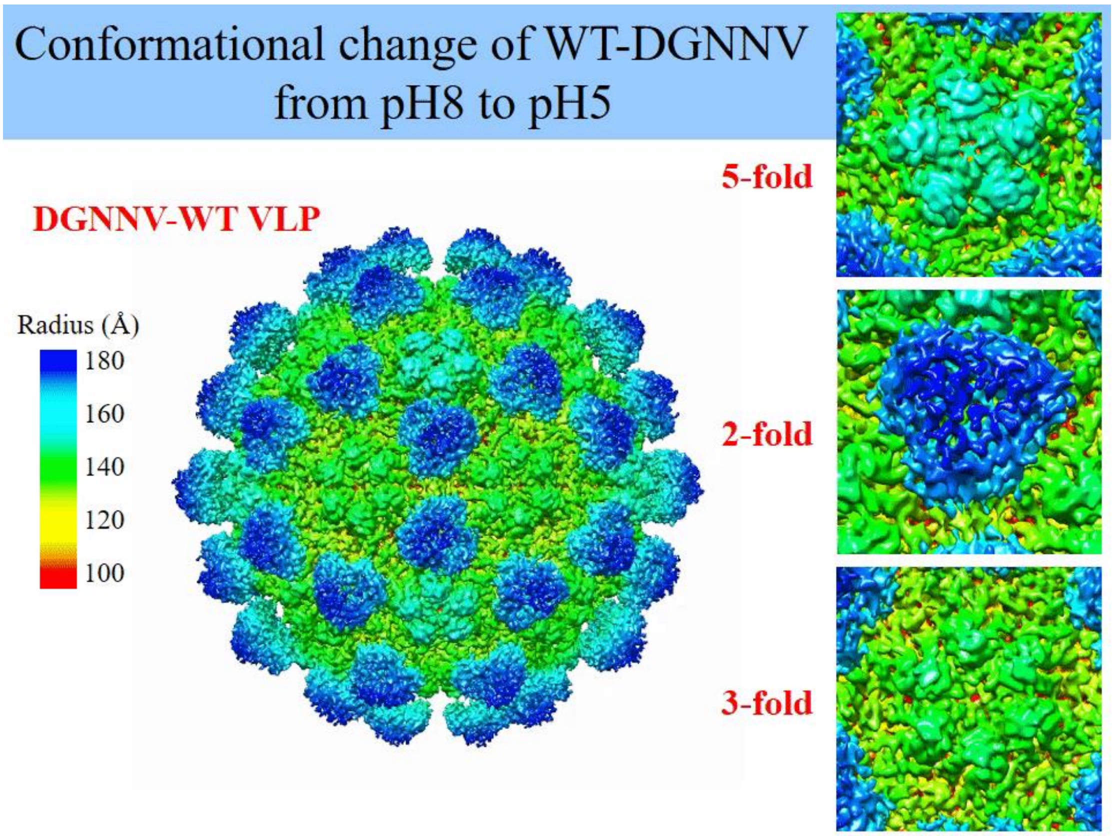

**Movie S2. Enlarged views of GNNV protrusion conformational change from pH 8.0 to 5.0.**

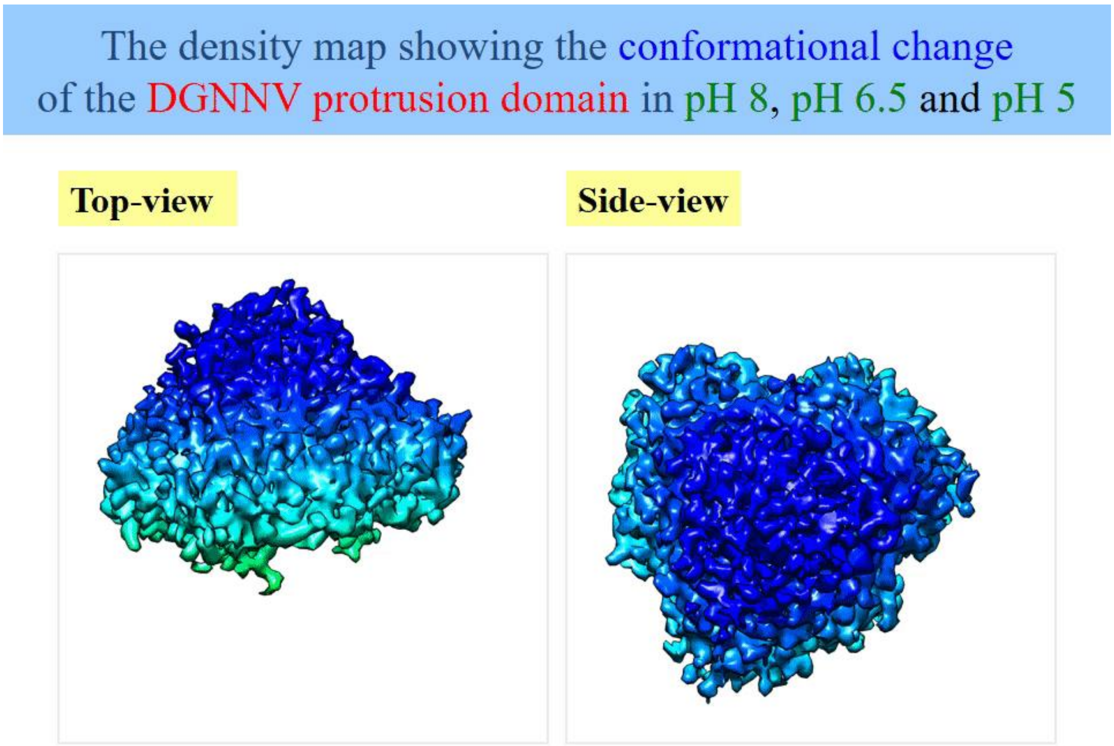

**Movie S3. Hypothetical atomic model of GNNV-P conformational change from pH 8.0 to 5.0.**

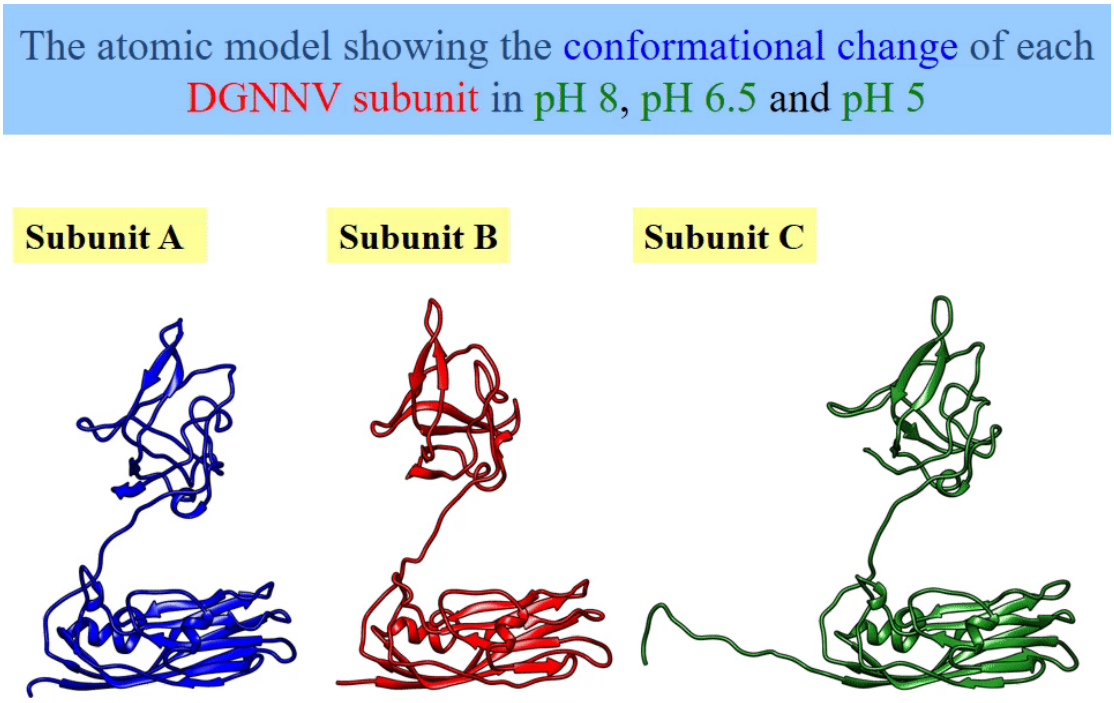
